# Dense Investigation of Variability in Affect (DIVA): A Neuroimaging Study of Premenopausal Female Participants

**DOI:** 10.1101/2024.04.16.589598

**Authors:** Katherine L. Bottenhorn, Taylor Salo, Julio A. Peraza, Michael C. Riedel, Jessica S. Flannery, Adam Kimbler, Alfredo Toll, Diego Suarez, Francis M. Cruz, Israel Zagales, Nayade Caldes, Olivia Dolan, Ruth Zagales, Matthew T. Sutherland, Robert W. Laird, Angela R. Laird

**Affiliations:** Department of Population and Public Health Sciences, University of Southern California; Department of Psychology, Florida International University; Lifespan Informatics and Neuroimaging Center (PennLINC), Department of Psychiatry, Perelman School of Medicine, University of Pennsylvania, Philadelphia, PA, United States; Lifespan Brain Institute (LiBI) of Penn Medicine and CHOP; Department of Physics, Florida International University; LTI Engineering and Software; Department of Psychology and Neuroscience, University of North Carolina, Chapel Hill; University of Alabama at Birmingham Marnix E. Heersink School of Medicine; Indiana University School of Medicine

## Abstract

The rise of large neuroimaging datasets and multi-dataset mega-analyses brings the power to study interindividual differences in brain structure and function on a heretofore unseen scale. However, unknown and poorly characterized intra-individual variability continues to undermine the detection of robust brain-behavior associations and, ultimately, our understanding of the brain on the whole. Women’s and reproductive health underlie variability in more than half of the population, but have long been overlooked in the study of both inter- and intra-individual differences in the brain. To this end, the Dense Investigation of Variability in Affect (DIVA) Study was designed to study intra-individual variability in the brain and behavior across the menstrual cycle in a small cohort of premenopausal female participants. The DIVA Study acquired weekly actigraphy, self-report, biospecimen, and both functional and structural magnetic resonance imaging data with concurrent peripheral physiological recordings. These data facilitate the study of several common sources of variability in the brain and behavior: the menstrual cycle and ovarian hormones, sleep, stress, exercise, and exogenous sources of hemodynamic variability.

## Introduction

As human neuroimaging seeks to identify brain-phenotype associations, questions of variability and statistical power continue to arise. Most recently, Marek and colleagues assessed brain-phenotype associations across two large datasets (N > 1,000) and found that sample sizes needed for the statistical power to detect the small brain-phenotype associations are several orders of magnitude larger than those of typical neuroimaging studies (Marek et al., 2022). While these findings may raise concerns, they do not necessarily invalidate any study without the resources to collect data from thousands of individuals. One interpretation of these findings is that brain-phenotype associations in cross-sectional brain-wide association studies (BWAS) are artificially small because trait-relevant individual differences are swamped by within-individual or processing-related variability (Bandettini et al., 2022). Characterizing this within-individual variability is among the goals of precision neuroscience (Poldrack, 2017). Moving from sparse sampling schemes (i.e., few observations across many individuals) to dense sampling schemes (i.e., many observations across fewer individuals) can provide greater characterization of both within- and between-individual variability (Gratton et al., 2022; Naselaris et al., 2021). Dense, longitudinal designs have already contributed quantitative estimates of sources of variability in functional connectivity (Gratton et al., 2018), of anatomical variability in large-scale functional networks (Seitzman et al., 2019), of weather-induced variability in functional neuroimaging (Di et al., 2022), of the impacts of food and caffeine consumption on functional brain networks (Poldrack et al., 2015), and of endocrine influences on functional brain networks (Mueller et al., 2021; Pritschet et al., 2020, 2021). Updates to imaging sequences and experimental design build on this increased precision, by mitigating time series variance for clearer identification of meaningful individual differences (Elliott et al., 2021). Specifically, multi-echo functional magnetic resonance imaging (fMRI) offers significant improvements in mitigating effects of noise (DuPre et al., 2021; Kundu et al., 2017; Lynch et al., 2021), as does concurrently collecting non-neural physiological measures (e.g., heart rate, respiration) (Caballero-Gaudes & Reynolds, 2017; Glover et al., 2000). Furthermore, naturalistic stimuli provide more reliable estimates of functional network connectivity (Wang et al., 2017) and greater ecological validity than resting-state or traditional task paradigms, while simultaneously engaging multimodal sensory processing, attention, and multiple aspects of cognition (Bottenhorn et al., 2018). Dense, longitudinal neuroimaging studies provide the data necessary to quantify variability in the brain and to identify sources of variability contributing to and confounding brain-phenotype associations, but there remain several open questions regarding the characterizing these sources of variability.

These open questions include the variable nature of the hemodynamic blood-oxygen level-dependent (BOLD) response and the impacts of data processing and non-neural physiological noise on estimates of brain function and functional connectivity from fMRI. Variability in the hemodynamic response (HR) was first acknowledged more than 20 years ago (Aguirre et al., 1998), exists both between individuals and within individuals, across regions of the cortex and across physiological states (Aguirre et al., 1998; Buckner et al., 1998; Handwerker et al., 2004, 2012). Within-individual variability in HR across the brain has been associated with proximity to large blood vessels, but is rarely incorporated into fMRI data analysis. Furthermore, ingestion of caffeine and over-the-counter pain and fever reducers that inhibit cyclooxygenase (e.g., ibuprofen) has been linked to changes in the hemodynamic response (Handwerker et al., 2012; Liu et al., 2004).

Despite decades of research regarding endocrine influences on the brain in non-human animals, the role of the brain as a crucial node of the endocrine system, and the presence of hormone receptors across the brain, and the role of steroid hormones as neurotransmitter agonists, human neuroimaging research concerning endocrine influences on the brain is limited. A large and notable source of neuroendocrine dynamics is the menstrual cycle, characterized by 8-fold changes in estradiol and 80-fold changes in progesterone over 24 to 34 days (Bull et al., 2019; Stricker et al., 2006). However, of human neuroimaging studies that directly measure hormones, fewer than 8% include more than 3 time points per individual and only 30% include more than 2 time points (reviewed in (Dubol et al., 2021)). Many such studies focus on comparing two phases of the menstrual cycle, which are defined by uterine and ovarian physiological changes that are accompanied by hormonal changes. As estradiol and progesterone fluctuations across the menstrual cycle are large, curvilinear, and vary greatly between individuals (Fehring et al., 2006), these sampling designs provide a poor estimation of neuroendocrine dynamics and are otherwise of relatively low quality (Dubol et al., 2021). While the literature to date has uncovered both structural and functional changes associated with hormone fluctuations over the course of the menstrual cycle, the experimental and sampling designs used in most of these studies impart nontrivial bias.

Here, we describe a dense, longitudinal study incorporating endocrine, physiological, multimodal neuroimaging, actigraphy, and behavioral data to investigate within-individual variability in the brain across the menstrual cycle. These data have already been used to test strategies for mitigating MR-related noise in peripheral electrophysiological data acquired concurrently with multi-band and multiecho fMRI sequences (Bottenhorn et al., 2021) and to identify contraceptive-related functional connectivity via predictive connectomics, in combination with data from the 28andMe and 28andOC studies (Bottenhorn et al., forthcoming; Pritschet et al., 2020).

The goal of the DIVA Study was to assess variability in several aspects of brain structure and function associated with endocrine fluctuations across the menstrual cycle and with hormonal contraception (HC). This includes variability in the hemodynamic response function, brain structure, brain function, and functional brain connectivity and contributions of lifestyle factors, affective and behavioral factors, and cognitive contexts. The DIVA Study recruited three premenopausal female participants (one naturally cycling, two using HC). Participants wore activity trackers throughout the duration of the study, completed weekly MRI scanning sessions with concurrent physiological recordings, and semiweekly collection of saliva samples and self-report behavioral measures. The imaging protocol included a rich battery of functional and structural scans. To maximize sampling across different phases of the menstrual cycle, the imaging protocol varied per scanning session. These data were collected to assess several common sources of variability in the brain and behavior that are frequently overlooked in human neuroimaging studies: the menstrual cycle and ovarian hormones, sleep, stress, exercise, and exogenous sources of hemodynamic variability.

## Methods

### Participants

This study included two pilot participants and three primary participants. While the original conception of DIVA was to collect data spanning three complete menstrual cycles per participant, with functional imaging tasks balanced between menstrual cycle phases, data collection was interrupted and ultimately truncated in March 2020 due to the global COVID-19 pandemic.

Pilot data were collected from two premenopausal, female participants (ages 25 and 40 years). Hormonal contraceptive use information was not collected from these individuals, as they did not undergo repeated scanning or saliva collection for hormone assessments.

Data for the primary study were collected from three premenopausal, female participants (“Blossom”, “Bubbles”, and “Buttercup”; age range = 26-31 years). At the time of data collection, two participants were using hormonal contraceptives (Blossom: 0.035 mg ethinyl-estradiol, 0.025 mg norgestimate, Feymor, Amneal Pharmaceuticals; Buttercup: 0.02 mg ethinyl-estradiol, 1 mg norethindrone acetate, Blisovi Fe). The third (Blossom) was freely cycling, with a history of regular menstrual cycles, who had not used hormonal contraceptives in the prior year. Participants completed behavioral assessments and collected saliva samples twice a week, 3-4 days apart, completed MRI scanning sessions once a week (on a behavioral & hormone collection day), and wore a FitBit activity tracker for the duration.

Written, informed consent was obtained from each participant before data collection began, in accordance with Florida International University’s Institutional Review Board approval.

### Activity tracking

Participants wore FitBit Charge HR 2 wearable devices to track activity, heart rate, and sleep patterns over the course of the study. The FitBits worn by participants in this study combine accelerometry and optical heart rate monitoring, at a 1 Hz sampling rate, to provide information about the wearer’s physical activity and the quality and duration of their sleep. They have been validated against polysomnography, in addition to research-grade accelerometers, and electrocardiograms, and indirect calorimetry (Bagot et al., 2018; Diaz et al., 2015; Mantua et al., 2016).

### Self-report measures

All self-report measures were acquired on a web browser on the participant’s personal device, via Qualtrics XM online surveys. These included trait measures, collected once, and state measures, collected twice a week, shortly after saliva sample collection, throughout the duration of the study.

#### Trait measures

Prior to the first visit for each of the DIVA participants, three trait measures were collected: the Behavioral Inhibition System/Behavioral Activation System (Carver & White, 1994), to assess individual tendencies toward appetitive or aversive motivations in their behavior; the Multi-Gender Identity Questionnaire (Joel et al., 2014), to assess the perception of gender identity; and the Mathematics Anxiety Rating Scale (Alexander & Martray, 1989), to assess a range of specific tensions and apprehensions associated with learning and being tested on mathematics.

#### State measures

For tracking factors associated with BOLD signal and affective variability, state measures were assessed semiweekly. These include the Pittsburgh Sleep Quality Index (Buysse et al., 1989), adapted to assess sleep quality on a weekly basis; the expanded form of the Positive and Negative Affect Schedule (Watson & Clark, 1999), to assess affective emotional states contributing to positive and negative emotional experiences; the Perceived Stress Scale (Cohen et al., 1994), adapted to assess feelings and thoughts concerning stressful events over the past week; and the Godin Leisure-Time Exercise Questionnaire (Godin & Shephard, 1985), to assess the amount of time spent doing vigorous, moderate, and leisurely exercise on a weekly basis. In addition, a physical state questionnaire was administered to assess menstrual cycle duration and recent birth control, caffeine, nicotine, acetaminophen, ibuprofen, and aspirin use.

Finally, following MRI scanning sessions, participants completed a post-scan questionnaire to assess whether participants fell asleep during the MRI scan, their perceived effort on each in-scanner task, and their attitudes toward characters in the episodes of Stranger Things that were viewed in the scanner.

### Hormone data

Endocrine measures include salivary estradiol, progesterone, and cortisol concentrations. Saliva samples were collected via passive drool into 2 mL sterile cryovials shortly after waking twice a week (3-4 days apart). Participants reported the time at which the sample was collected as a part of the larger self-report battery. Samples were stored at -20 C until shipping, following completion of the study, to Salimetrics’ SalivaLab (Carlsbad, CA). They were then assayed using the Salimetrics Salivary Estradiol Assay Kit (Cat. No. 1-3702) and the Salimetrics Salivary Progesterone Assay Kit (Cat. No. 1-1502), without modifications to the manufacturers’ protocols. All samples were assayed in duplicate and values reflect the average salivary concentration.

### Physiological data acquisition

Physiological data were acquired simultaneously with MRI data, using MR-compatible equipment from BIOPAC Systems, Inc.: electrocardiography (ECG) for heart rate, chest-belt recording for respiration, and electrodermal activity (EDA) for skin conductance. A BIOPAC MP150 system was connected to subject leads through the MRI patch panel with MRI-RFIF filters by two standard MEC-MRI cables that ran to the bore, without loops, and then ran parallel to the subject. Three radiotranslucent EL508 electrodes with GEL100 and 15 cm long LEAD108B leads were used to collect ECG recordings, together with an ECG100C-MRI amplifier. Electrodes were placed in a bipolar monitoring configuration: two electrodes placed 6-8 inches apart diagonally across the heart from left rib cage to the right clavicle, and the ground electrode was placed 6-8 inches away on the right rib cage. Radiotranslucent EL509 electrodes with GEL101 and LEAD108B leads were used to acquire EDA recordings, together with an EDA100C-MRI amplifier. Leads were placed on the thenar and hypothenar eminences of the palm of the participant’s non-dominant hand. A TSD221-MRI transducer and belt, placed snugly around the abdomen, were used to acquire respiration signal. Physiological data (i.e., ECG, EDA, and respiration) were acquired at a rate of 2000 Hz, throughout the duration of the scanning session: beginning when participants were loaded on the scanner bed and continuing until the scanner bed exited the bore at the end of the scanning session. All recordings include several minutes of data, per participant per session, collected in the absence of an MR pulse sequence.

Additionally, a trigger channel was included to record a binary trigger signal indicating whether an fMRI scan was being acquired or not. This channel was triggered by a signal originating from the task scripts on the stimulus computer, rather than a direct signal from the scanner.

### MRI data acquisition

MRI data were acquired on a 3T Siemens Prisma MRI scanner with a 32-channel head/neck coil at Florida International University (Miami, FL USA), using the VE11C software. Sequence parameters and file naming conventions are summarized in Table 1.

**Table 1.**
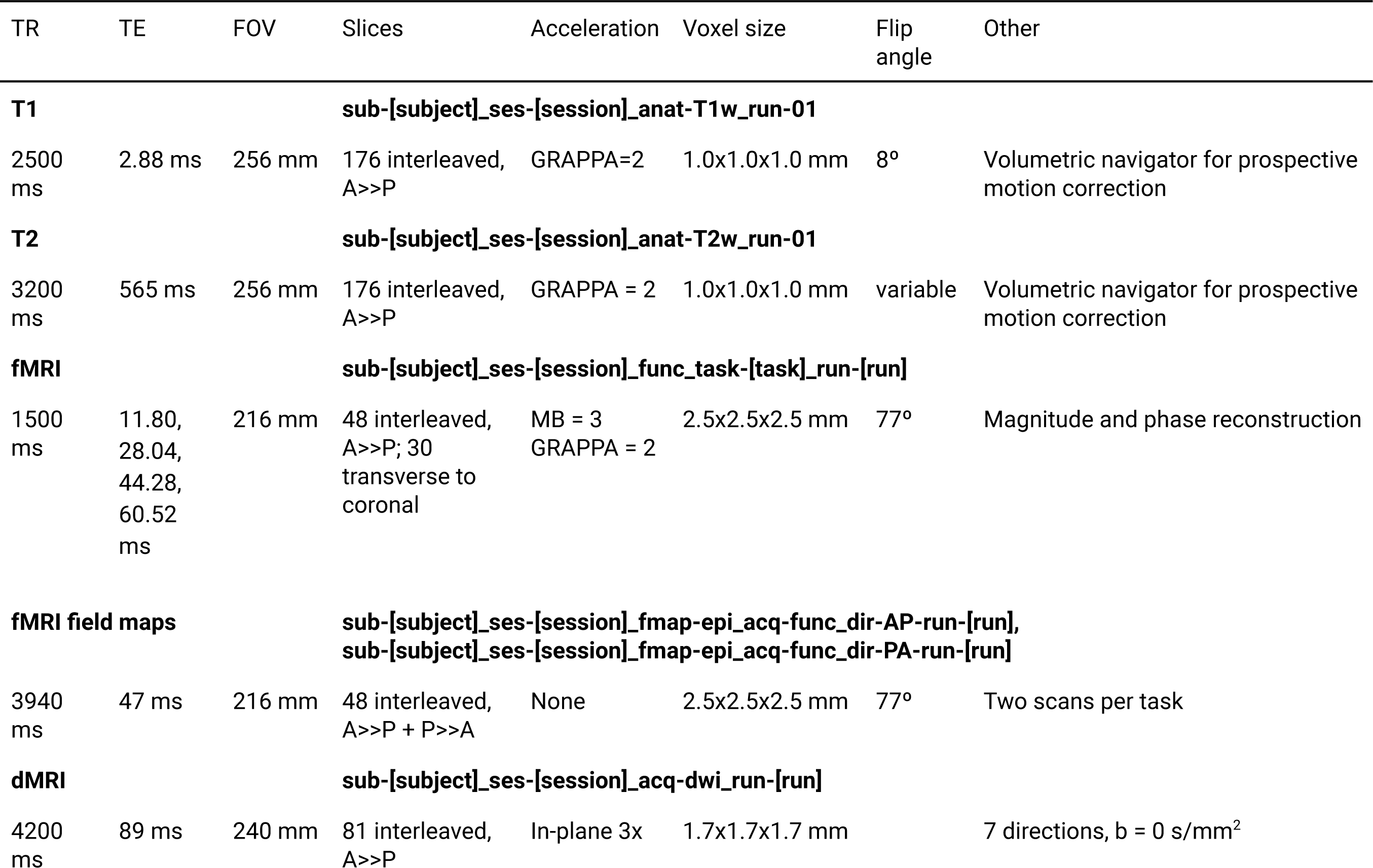
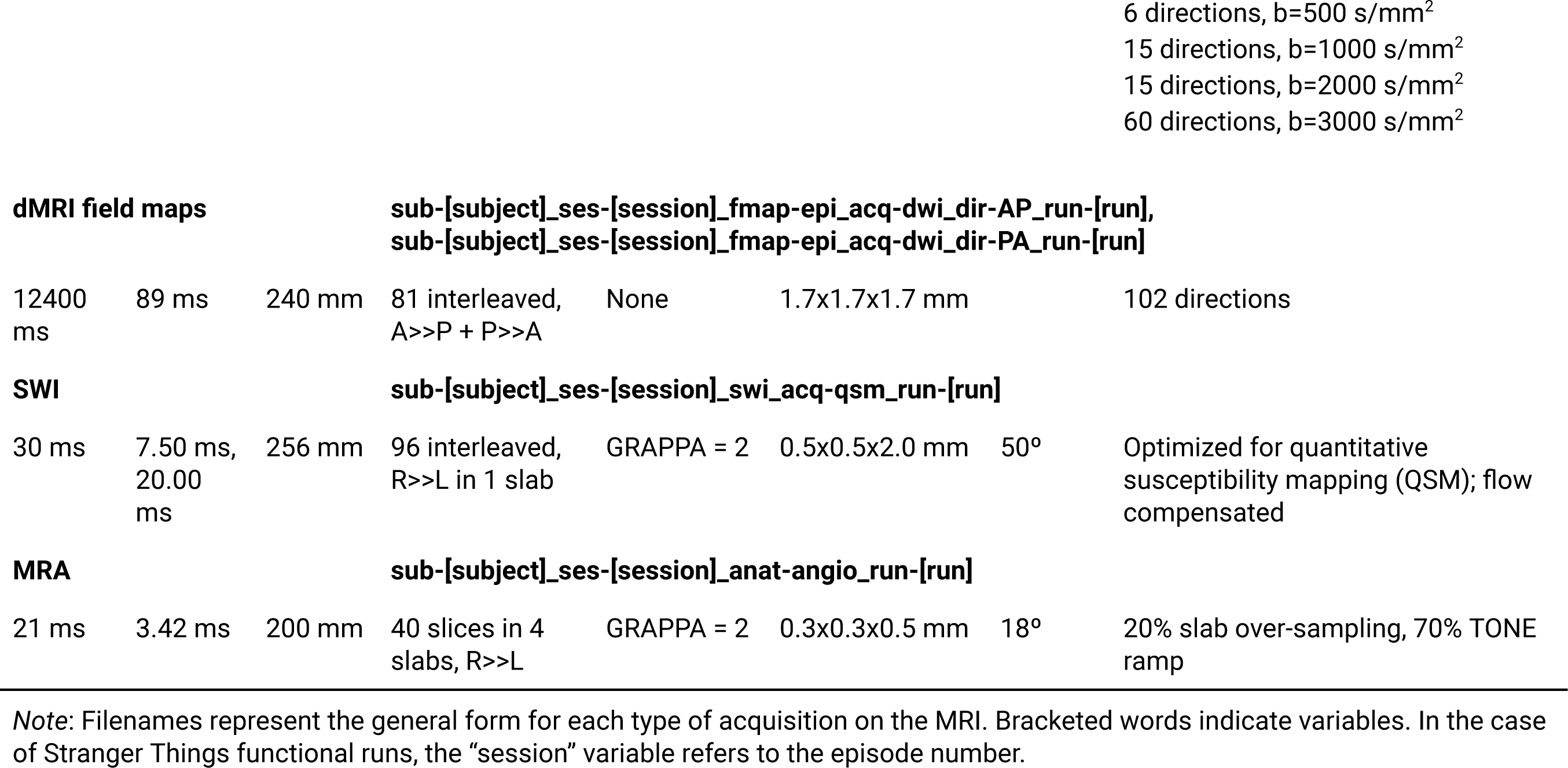
MRI sequence parameters.

Functional MRI scans were acquired with the CMRR multiband sequence (version 016a). Each functional run included four echoes (echo times, TEs=11.8, 28.04, 44.28, 60.52ms) and both magnitude and phase data reconstruction. The runs had the following parameters: repetition time, TR=1500 ms; multiband factor=3; flip angle, FA=77°; matrix size=86x86; voxel size=2.5x2.5x2.5 mm; field of view, FOV=216 mm; 48 slices acquired in interleaved ascending order, at a 30° transverse-to-coronal orientation.

Prior to each functional scan, two B_0_ calibration scans were acquired: one with phase encoding from anterior to posterior; the other, posterior to anterior. These scans had TR = 3940 ms, TE = 47 ms, FOV = 216 mm, 2.5 mm isotropic voxels, 48 interleaved slices, at a 30° transverse-to-coronal orientation.

Structural T1-weighted images were acquired using a 3D T1w inversion prepared RF-spoiled gradient echo scan, the same sequence used by the Adolescent Brain Cognitive Development℠ Study (ABCD Study®) (Casey et al., 2018), with anterior-to-posterior phase encoding direction, TR = 2500 ms, TE = 2.88 ms, TI = 1070 ms, FOV = 256 mm, in-plane acceleration (GRAPPA=2), and 1 mm^3^ isotropic voxels, with Volumetric Navigators (vNav) for prospective motion correction (Tisdall et al., 2012).

T2-weighted anatomical MRI scans 3D T2-weighted fast spin echo the same sequence used by the Adolescent Brain and Cognitive Development (ABCD) Study℠ (Casey et al., 2018), with anterior-to-posterior phase encoding direction, TR = 3200 ms, TE = 565 ms, FOV = 256 mm, in-plane acceleration (GRAPPA=2), and 1 mm^3^ isotropic voxels with variable flip angles and Volumetric Navigators (vNav) for prospective motion correction.

Diffusion-weighted MRI scans were acquired with the same high angular resolution diffusion imaging (HARDI) multiband EPI sequence, with integrated static field distortion correction, used by the ABCD Study (Casey et al., 2018; Hagler et al., 2019) that acquires 96 diffusion directions with 4 b-values (6 directions, b=500 s/mm^2^; 15 directions, b=1000 s/mm^2^; 15 directions, b=2000 s/mm^2^; and 60 directions, b=3000 s/mm^2^) and 7 b=0 volumes. The sequence had TR = 4200 ms, TE = 89 ms, FOV = 240 mm, 1.7x1.7x1.7 mm isotropic voxels, 81 interleaved slices acquired anterior-to-posterior, a multiband factor of 3, and in-plane acceleration (GRAPPA = 2).

Prior to each diffusion scan, two B_0_ calibration scans were acquired: one with phase encoding from anterior to posterior; the other, posterior to anterior. These scans had TR = 12400 ms, TE = 89 ms, FOV = 240 mm, 1.7x1.7x1.7 mm isotropic voxels, 81 interleaved slices, at a 30° transverse-to-coronal orientation.

Susceptibility-weighted MRI (SWI) scans were acquired for quantitative susceptibility mapping (QSM) with channel-level reconstruction, per recommendations from (Haacke et al., 2015): TR = 30 ms, FA = 15 degrees, TEs = 7.5, 20 ms, FOV = 256 mm, 0.5x0.5x2 mm voxels, 96 transverse, interleaved slices in one slab acquired with a right-to-left phase-encoding direction, and in-plane acceleration (GRAPPA = 2).

Magnetic resonance angiograms (MRA) were acquired using a multi-slab, ramped flip-angle time-of-flight (TOF) sequence with TR = 21 ms, TE = 3.42 ms, FOV = 200 mm, 0.3x0.3x0.5 mm anisotropic voxels, 4 slabs (40 slices/slab, GRAPPA = 2), 20% slab oversampling, an 18-degree flip angle and a 70% TONE ramp.

### Task design

Seven tasks were employed in the dataset: an A1/V1/M1 functional localizer, an arithmetic task, a dual self- and other-referential processing task, an emotion induction and regulation task, a probabilistic selection task, a resting-state task, and a film-viewing task.

#### Sensorimotor functional localizer

The functional localizer task comprises four conditions: motor, visual, combined motor/auditory, and combined visual/auditory across a block design run to prioritize detection and a rapid event-related design to prioritize BOLD response estimation in the target regions. Motor trials involve a text-based prompt to tap one’s fingers as quickly as possible. Visual trials involve a flashing checkerboard. Auditory trials involve the presentation of a randomly selected public domain song.

In the block design run, 14-second trial blocks are separated by 14-second inter-block fixation blocks. Condition order was randomized, but consecutive blocks of the same condition were not allowed.

In the event-related run, intertrial intervals were randomly drawn from a right-skewed Gumbel distribution with a mean of 4 seconds and a scale of 1 second. Resulting values were restricted to between 2 and 8 seconds, and were rounded to the nearest tenth of a second. Trial durations were drawn from a uniform distribution limited to the range of 0.5 to 4 seconds, and then rounded to the nearest tenth of a second. Condition order was randomized, but consecutive trials of the same condition were not allowed. There were 60 trials overall, with 15 trials of each condition.

The auditory/visual/motor localizer task was implemented in PsychoPy (Peirce et al., 2019). The task includes two 7 minute 30 second runs.

#### Arithmetic task

Trials consist of three stages: equation, comparison value, and feedback. In the equation stage, an equation is presented which participants must solve. This equation may be addition, subtraction, division, multiplication, or a baseline in which a single number is presented. In the comparison stage, a single value is presented and the participant must respond whether the solution to the previous equation is (1) less than, (2) equal to, or (3) greater than the comparison value. Finally, in the feedback stage, the participant is provided with feedback on their response. That feedback may be either informative, in which case a smiley face indicates that their response was correct and a frowny face indicates that their response was incorrect, or uninformative, in which case a neutral face is shown regardless of trial accuracy.

Trial difficulty varied based on operator type (addition, subtraction, division, multiplication, and baseline), value size (i.e., multiplying larger values is generally harder than smaller values), and the scale of the difference between the comparison value and the equation solution. Additionally, both the equations and comparison values were provided in either numerical or textual form.

The arithmetic task was implemented in PsychoPy. The task includes an out-of-scanner training run, as well as two in-scanner, 7 minute 30 second runs.

#### Dual Self- and Other-Referential Processing/Flanker task (SORPF)

The Dual Self- and Other-Referential Processing/Flanker (SORPF) Task was adapted from Alarcón and colleagues (Alarcón et al., 2018). This task combines the self and other referential processing task and the Eriksen flanker task in a block design, with four conditions: “self”, “other”, “malleable”, and “flanker”. In the “self” condition, the participant views an image of themself paired with a descriptive word or phrase and is asked “does this word describe you?” to which they are prompted to respond either “yes” or “no”. This condition is designed to engage self-referential processing. In the “other” condition, the participant views an image of a familiar person (in this case, a character from the TV show Stranger Things) paired with a descriptive word or phrase and is asked “does this word describe the person shown?”, to which they are prompted to respond either “yes” or “no”. This condition is designed to engage other-referential processing. In the “malleable” condition, the participant views an image of a stranger paired with a descriptive word or phrase. The participant is then asked, “Can this change?”, and prompted to respond “yes” or “no”, depending on whether the descriptive word is something that can change about a person. This condition is included as a high-level control, perceptually and temporally matched to the “self” and “other” conditions. Finally, in the “flanker” condition the participant performs 5 randomized trials of the Eriksen flanker task. Each trial is 800 ms, during which the participant is instructed to indicate the direction that the center arrow is pointing as quickly as possible. Trials are randomized to include congruent trials (center arrow points in the same direction as the flanking arrows) and incongruent trials (center arrow is pointing in the opposite direction as the flanking arrows). “Flanker” conditions are interspersed between each of the previously described conditions as a mental palate cleanser, in the form of an attentionally-demanding, out-of-domain task, to interrupt ongoing cognition in an attempt to separate self-from other-referential processing. This task was implemented in EPrime and comprises two 7 minute 30 second runs.

#### Emotion Induction and Regulation Task (EIRT)

The Emotion Induction and Regulation Task (EIRT) is a fast, event-related task adapted from Blair and colleagues (Blair et al., 2012), and similar to that of Ochsner et al. (Ochsner et al., 2004). The task uses negative and neutral images from the International Affective Picture System (IAPS) (Lang et al., 1997)and the Self-Assessment Manikin (SAM) instrument (Betella & Verschure, 2016) to assess emotion induction and regulation with two different instructions: VIEW (induction; neutral and negative images) and BETTER (regulation, reappraisal; negative images only). There are fifteen neutral and fifteen negative images each, and each trial randomly pairs an image with an appropriate instruction, followed by the valence SAM scale during which they are asked to rate the valence of the image on a scale from 1 (most negative) to 4 (most positive). Participants completed two runs, with a total of 30 10-second trials per run, interspersed with jittered fixations between each trial ranging from 500 ms to 1500 ms. The EIRT was implemented in E-Prime and comprises two 7 minute 30 second runs.

#### Probabilistic Selection Task (PST)

The Probabilistic Selection task (PST) is a reinforcement learning task used to separately calculate positive and negative reinforcement learning rates and learning performances, in addition to behavioral sensitivity to positive and negative feedback via win-stay and loose-shift behavioral choices (Frank et al., 2004).

The out-of-scanner practice run was completed the first time the participant was introduced to the task. This practice included an initial instruction phase (∼4 min) in which the task was explained to the participant and six example trials. This was followed by a practice phase (∼5 min) in which the participant practiced all task procedures (30 training trials and 10 testing trials) with different stimuli than those presented in the actual task as to prevent any pre-task learning about the stimuli. Once the experimenter ensured that the participant understood the task rules, the participant completed the real task during their MRI scans.

During the in-scanner training run, participants were presented with 3 different stimuli pairs (AB, CD, EF) and learned, through ‘trial and error’, to choose which stimulus was ‘the best choice’ based on probabilistic feedback indicating correct or erroneous selections. The training run consisted of 60 trials. Each trial started with a choice screen (2500 ms) in which participants were presented with 1 of the 3 stimuli pairs. Stimuli in this task were affectively-neutral, nonrepresentational white markings/symbols on black background that have been used in previous implementations (Frank et al., 2004). The side of the screen each stimulus was displayed on changed each trial, in a random sequence. Participants choose one of the stimuli using a button response box. They were then presented with probabilistic feedback (1000 ms) consisting of a green smiling emoji for correct feedback and a red frowning emoji for erroneous feedback. In AB trials, stimulus A leads to positive feedback 80% of the time whereas stimulus B leads to negative feedback 80% of the time. CD and EF pairs are less reliable, such that stimulus C leads to positive feedback on 70% of selections and D leads to negative feedback on 70% of selections. Stimulus E leads to positive feedback on 60% of EF trials and F leads to negative feedback on 60% of EF trials. Over the course of training, participants learn to choose stimuli A, C, and E more often than B, D, or F. The probabilistic properties of symbols are always randomly reassigned at the start of the task (e.g. which symbol will be the “A” stimulus) to prevent any symbol-specific effects. A fixation cross is presented between each trial with a jittered duration between 1000 and 3200 ms.

During the in-scanner testing run, novel combinations of stimuli pairs that included either an A (AC, AD, AE, AF) or a B (BC, BD, BE, BF) were presented (2500 ms) and no feedback was provided. Again, a fixation cross with a jittered duration was presented between each trial.

The PST was implemented in E-Prime and includes an out-of-scanner, experimenter-guided practice run, as well as two in-scanner runs, a training run (5 minutes 54 seconds) followed by a testing run (8 minutes 40 seconds).

#### Resting-state

The resting-state task was implemented in E-Prime. The task was 7 minutes and 30 seconds long, in which participants were instructed to keep their eyes open and focused on a fixation cross in the center of the screen. Participants completed 1 to 4 runs, depending on the session.

#### Film-viewing task

The film-viewing task was implemented in PsychoPy. Participants were presented in each session with one episode from season one of the Netflix series Stranger Things (Duffer & Duffer, 2017). The episodes were broken up into 6-7 runs ranging in length from 5 minutes 35 seconds to 10 minutes 54 seconds.

#### Stranger Things Annotations

To facilitate analysis of the naturalistic film-viewing fMRI data acquired while participants watched the television show Stranger Things, the visual features, emotional valence, and emotional arousal of each episode were annotated with TR resolution. Briefly, each run of each episode was broken into TR-length clips (i.e., 1.5 second clips). Authors AT, DS, FC, IZ, NC, OD, and RZ viewed each clip repeatedly using the VLC Media Player (https://www.videolan.org) and denoted the most salient features, in nouns and verbs, using words and senses from the WordNet database (https://wordnet.princeton.edu). WordNet is a lexical database that groups nouns, verbs, adjectives, and adverbs in the English language into “SynSets” of cognitive synonyms which are linked by conceptual, semantic, and lexical relations. Annotators also noted the presence of any text and major characters on the screen during each TR-clip. Additionally, each TR-clip was annotated according to the emotional valence and arousal of the clip, using the Self-Assessment Manikin (Betella & Verschure, 2016; Bradley & Lang, 1994) on scales from 1 (most negative emotion, lowest arousal) to 7 (most positive emotion, highest arousal).

### Data processing

All of the in-house scripts used to organize and process this data are available at https://github.com/NBCLab/diva-project. External tools and packages are referenced and linked throughout.

#### Self-report

Each measure was scored according to its published guidelines. Trait measures, administered once to each participant, were summarized across participants by their means and standard deviations. State measures, administered biweekly to each participant, were summarized per participant, across sessions, by their means and standard deviations.

#### Hormone data

Assays per hormone were performed in duplicate by Salimetrics and summarized by averaging the two results per session, then plotted per participant across sessions. Measurements below the lower limit of detection were excluded.

#### Physiological data

Physiological recordings (i.e., ECG, EDA, respiration) were converted to BIDS-compatible format and organized using a customized clone of phys2bids (phys2bids.readthedocs.io; https://github.com/tsalo/phys2bids/releases/tag/diva-paper) specific to the physiological data acquisition setup used for this dataset.

From the raw ECG recordings, heart rate was calculated and the signal quality was estimated from the signal’s kurtosis and an automated, composite heuristic (Zhao & Zhang, 2018). Kurtosis, or the fourth moment of the ECG signal in the time domain, has been used previously to estimate signal quality (del Río et al., 2011; Rahman et al., 2022), such that higher kurtosis indicates better quality. The heuristic quality index (or, “Zhao heuristic”) indicates whether an ECG recording is “unacceptable”, “barely acceptable”, or “excellent” by estimating four previously-validated indices of ECG signal quality (R peak detection, QRS wave power spectrum distribution, kurtosis, and baseline relative power) and using a fuzzy comprehensive assessment to categorize signal quality.

Objective measures of EDA recording quality depend on acquisition and analytic context. Here, we provide a visual example of pre-scan EDA recording and an EDA recording during a BOLD EPI scan (Figure 1, right column, rows 1 vs. 2).

**Figure 1.**
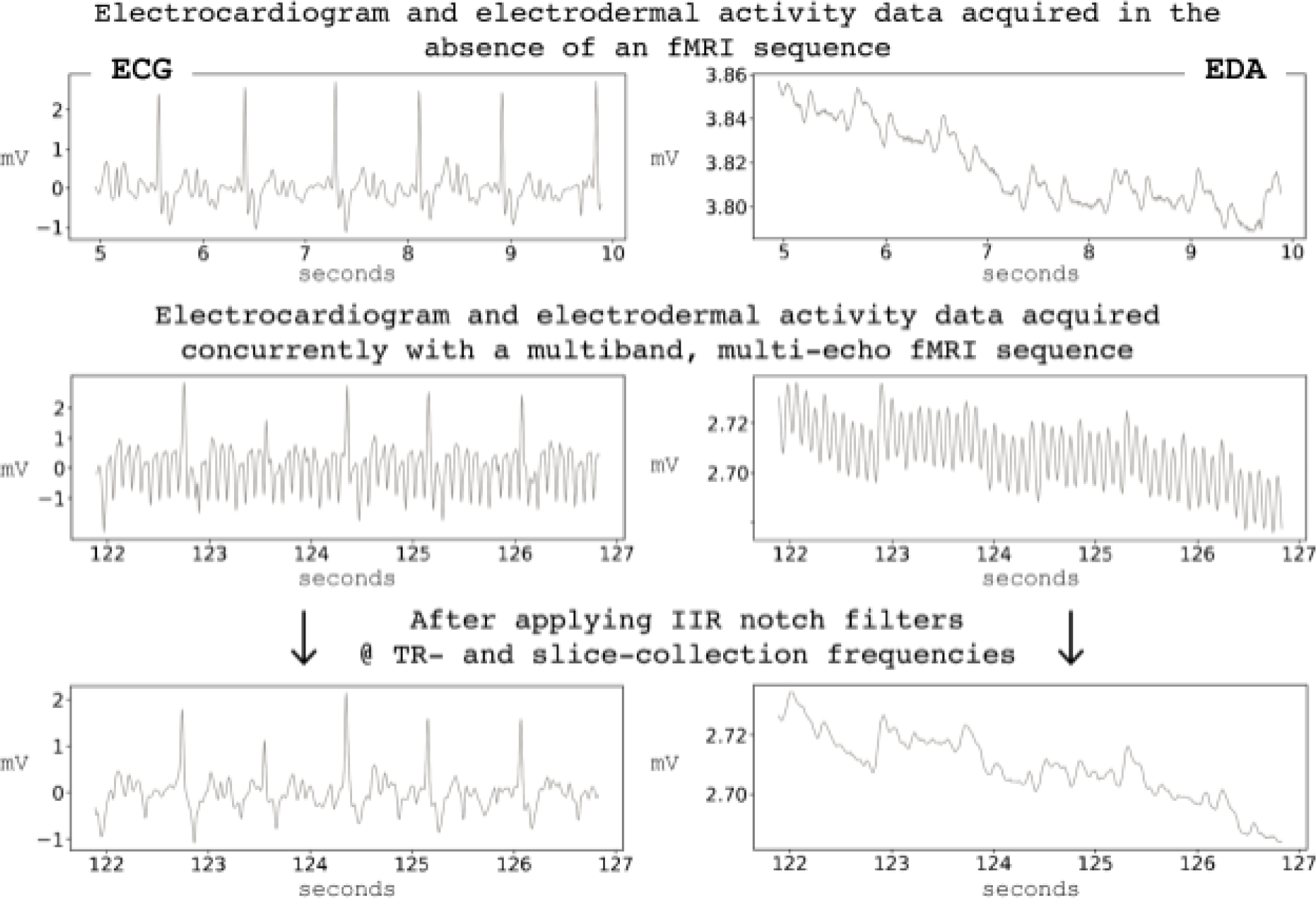
Example physiological signals (i.e., ECG, right; EDA, left) before (top row) and during (middle row) multiband, multi-echo fMRI sequence, and MR-specific filtering (bottom row).

Then, MR-related artifacts were removed from ECG and EDA recordings using physioComb (Bottenhorn, 2022; Bottenhorn et al., 2021) by applying notch filters centered at the slice collection frequency (i.e., number of slices / MB factor / TR; 10.67Hz) and the TR frequency (i.e., 1/TR; 0.67Hz), and their harmonics up to the Nyquist frequency (Figure 1, row 3). Finally, the data were preprocessed with a 0.5Hz low-pass filter, then by powerline filtering, following recommendations from NeuroKit2 (Makowski et al., 2021). The aforementioned quality metrics were recalculated following filtering and preprocessing (Figure 3).

**Figure 2.**
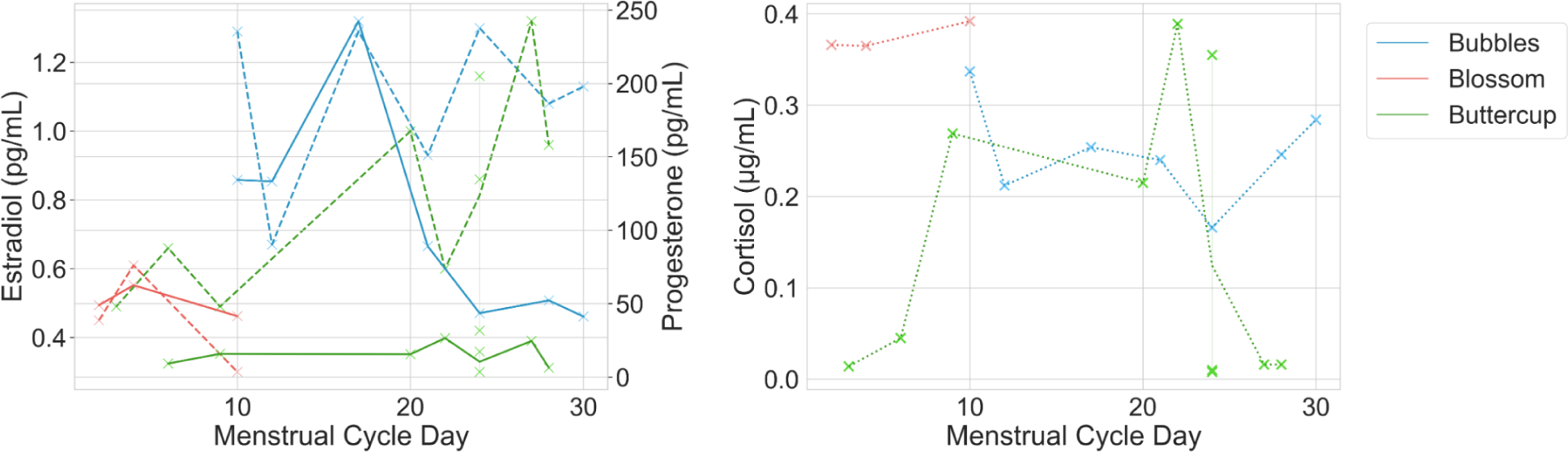
Salivary hormone concentrations throughout data collection, plotted by menstrual cycle day. Left: ovarian hormones estradiol (dashed) and progesterone (solid); right: cortisol (dotted). Note: Blossom and Bubbles use hormonal contraception, Buttercup is naturally cycling.

**Figure 3.**
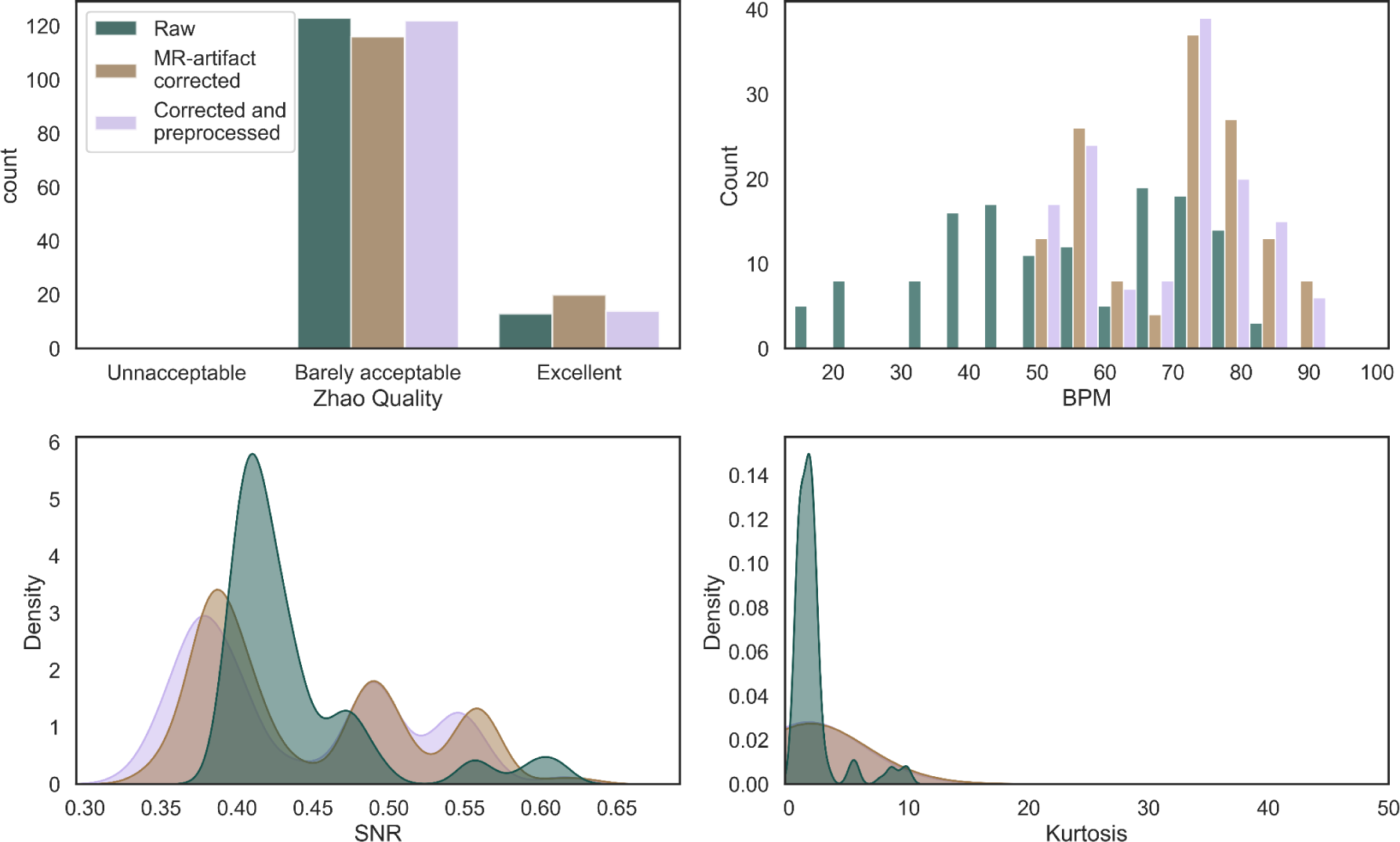
Average heart rate and electrocardiogram signal quality indices across the dataset.

Respiratory recordings were low-pass filtered at 3Hz and then respiration rate was calculated by identifying peaks (representing the transition from inhaling to exhaling) and then dividing 60 seconds/minute by the associated inter-breath interval in seconds.

### MRI data

#### MRIQC and fMRIPrep

Imaging data were converted to NiFTI images from DICOMs using dcm2niix (https://github.com/rordenlab/dcm2niix; v1.0.20200331). Then, MRIQC (mriqc.readthedocs.io; v22.0.6 and fMRIPrep (fmriprep.org; v22.0.0 were used to assess the quality of and preprocess T1, T2, and fMRI data (Esteban, Birman, et al., 2017; Esteban et al., 2019).

Briefly, MRIQC performs skull stripping, calculates a head mask, uses ANTS to normalize images to an MNI template brain, calculates an air mask under the base of the brain, and uses FSL’s automated segmentation tool (FAST) to segment tissue into white matter, gray matter, and cerebral spinal fluid. Then, image quality metrics (IQMs) are extracted from each image and reports are generated per participant and summarized across the dataset.

For each of the 3 BOLD runs found per subject (across all tasks and sessions), the following preprocessing was performed. First, a reference volume and its skull-stripped version were generated by aligning and averaging the first echo of 4 single-band references (SBRefs). A B0-nonuniformity map (or fieldmap) was estimated based on two (or more) echo-planar imaging (EPI) references with opposing phase-encoding directions, with 3dQwarp Cox and Hyde (1997) (AFNI 20160207). Based on the estimated susceptibility distortion, a corrected EPI (echo-planar imaging) reference was calculated for a more accurate co-registration with the anatomical reference. The BOLD reference was then co-registered to the T1w reference using bbregister (FreeSurfer) which implements boundary-based registration (Greve and Fischl 2009). Co-registration was configured with six degrees of freedom. Head-motion parameters with respect to the BOLD reference (transformation matrices, and six corresponding rotation and translation parameters) are estimated before any spatiotemporal filtering using mcflirt (FSL 5.0.9, Jenkinson et al. 2002). BOLD runs were slice-time corrected using 3dTshift from AFNI 20160207 (Cox and Hyde 1997, RRID:SCR_005927). The BOLD time-series (including slice-timing correction when applied) were resampled onto their original, native space by applying a single, composite transform to correct for head-motion and susceptibility distortions. These resampled BOLD time-series will be referred to as preprocessed BOLD in original space, or just preprocessed BOLD. A T2* map was estimated from the preprocessed BOLD by fitting to a monoexponential signal decay model with nonlinear regression, using T2*/S0 estimates from a log-linear regression fit as initial values. For each voxel, the maximal number of echoes with reliable signal in that voxel were used to fit the model. The calculated T2* map was then used to optimally combine preprocessed BOLD across echoes following the method described in (Posse et al. 1999). The optimally combined time series was carried forward as the preprocessed BOLD. First, a reference volume and its skull-stripped version were generated using a custom methodology of fMRIPrep. The BOLD time-series were resampled onto the following surfaces (FreeSurfer reconstruction nomenclature): fsnative, fsaverage5. The BOLD time-series were resampled into standard space, generating a preprocessed BOLD run in MNI152NLin2009cAsym space. First, a reference volume and its skull-stripped version were generated using a custom methodology of fMRIPrep. Several confounding time-series were calculated based on the preprocessed BOLD: framewise displacement (FD), DVARS and three region-wise global signals. FD was computed using two formulations following Power (absolute sum of relative motions, Power et al. (2014)) and Jenkinson (relative root mean square displacement between affines, Jenkinson et al. (2002)). FD and DVARS are calculated for each functional run, both using their implementations in Nipype (following the definitions by Power et al. 2014). The three global signals are extracted within the CSF, the WM, and the whole-brain masks. Additionally, a set of physiological regressors were extracted to allow for component-based noise correction (CompCor, Behzadi et al. 2007). Principal components are estimated after high-pass filtering the preprocessed BOLD time-series (using a discrete cosine filter with 128s cut-off) for the two CompCor variants: temporal (tCompCor) and anatomical (aCompCor). tCompCor components are then calculated from the top 2% variable voxels within the brain mask. For aCompCor, three probabilistic masks (CSF, WM and combined CSF+WM) are generated in anatomical space. The implementation differs from that of Behzadi et al. in that instead of eroding the masks by 2 pixels on BOLD space, the aCompCor masks are subtracted from a mask of pixels that likely contain a volume fraction of GM. This mask is obtained by dilating a GM mask extracted from the FreeSurfer’s aseg segmentation, and it ensures components are not extracted from voxels containing a minimal fraction of GM. Finally, these masks are resampled into BOLD space and binarized by thresholding at 0.99 (as in the original implementation). Components are also calculated separately within the WM and CSF masks. For each CompCor decomposition, the k components with the largest singular values are retained, such that the retained components’ time series are sufficient to explain 50 percent of variance across the nuisance mask (CSF, WM, combined, or temporal). The remaining components are dropped from consideration. The head-motion estimates calculated in the correction step were also placed within the corresponding confounds file. The confound time series derived from head motion estimates and global signals were expanded with the inclusion of temporal derivatives and quadratic terms for each (Satterthwaite et al. 2013). Frames that exceeded a threshold of 0.5 mm FD or 1.5 standardised DVARS were annotated as motion outliers. All resamplings can be performed with a single interpolation step by composing all the pertinent transformations (i.e. head-motion transform matrices, susceptibility distortion correction when available, and co-registrations to anatomical and output spaces). Gridded (volumetric) resamplings were performed using antsApplyTransforms (ANTs), configured with Lanczos interpolation to minimize the smoothing effects of other kernels (Lanczos 1964). Non-gridded (surface) resamplings were performed using mri_vol2surf (FreeSurfer).

Many internal operations of fMRIPrep use Nilearn 0.6.2 (Abraham et al. 2014, RRID:SCR_001362), mostly within the functional processing workflow. For more details of the pipeline, see the section corresponding to workflows in fMRIPrep’s documentation.

#### QSIPrep

Preprocessing was performed using *QSIPrep* 0.16.1 (Cieslak et al., 2021), which is based on *Nipype* 1.8.5 ((Gorgolewski et al., 2011, 2018); RRID:SCR_002502) and uses the FreeSurfer derivatives from fMRIPrep (above).

#### Diffusion data preprocessing

Any images with a b-value less than 100 s/mm^2 were treated as a *b*=0 image. MP-PCA denoising as implemented in MRtrix3’s dwidenoise (Veraart et al., 2016) was applied with a 5-voxel window. After MP-PCA, B1 field inhomogeneity was corrected using dwibiascorrect from MRtrix3 with the N4 algorithm (Tustison et al., 2010). After B1 bias correction, the mean intensity of the DWI series was adjusted so all the mean intensity of the b=0 images matched across each separate DWI scanning sequence.

FSL (version 6.0.5.1:57b01774)’s eddy was used for head motion correction and Eddy current correction (Andersson et al., 2016). Eddy was configured with a *q*-space smoothing factor of 10, a total of 5 iterations, and 1000 voxels used to estimate hyperparameters. A linear first level model and a linear second level model were used to characterize Eddy current-related spatial distortion. *q*-space coordinates were forcefully assigned to shells. Field offset was attempted to be separated from subject movement. Shells were aligned post-eddy. Eddy’s outlier replacement was run (ibid). Data were grouped by slice, only including values from slices determined to contain at least 250 intracerebral voxels. Groups deviating by more than 4 standard deviations from the prediction had their data replaced with imputed values. Data was collected with reversed phase-encode blips, resulting in pairs of images with distortions going in opposite directions. Here, b=0 reference images with reversed phase encoding directions were used along with an equal number of b=0 images extracted from the DWI scans. From these pairs the susceptibility-induced off-resonance field was estimated using a method similar to that described in (Andersson et al., 2003). The fieldmaps were ultimately incorporated into the Eddy current and head motion correction interpolation. Final interpolation was performed using the jac method.

Several confounding time-series were calculated based on the preprocessed DWI: framewise displacement (FD) using the implementation in *Nipype* (following the definitions by (Power et al., 2014)). The head-motion estimates calculated in the correction step were also placed within the corresponding confounds file. Slicewise cross correlation was also calculated. The DWI time-series were resampled to ACPC, generating a *preprocessed DWI run in ACPC space* with 2mm isotropic voxels.

Many internal operations of *QSIPrep* use *Nilearn* 0.9.2 ((Abraham et al., 2014), RRID:SCR_001362) and *Dipy* (Garyfallidis et al., 2014). For more details of the pipeline, see <u>the</u> <u>section corresponding to workflows in *QSIPrep*’s documentation</u>.

## Data Records

Data are shared on OpenNeuro (https://openneuro.org/datasets/ds002278; Salo et al., 2019), following organization and file conventions of the Brain Imaging Data Structure (Gorgolewski et al., 2016), along with a descriptive README file. All personally identifying information has been removed from these records. All scripts used to perform the processing and analysis presented in this manuscript are available on GitHub (https://github.com/NBCLab/diva-project). Table 1 includes the naming conventions for the imaging data and Table 2 provides information regarding the scans collected per participant, along with the number of scanning sessions across the menstrual cycle or oral contraceptive pill.

**Table 2.**
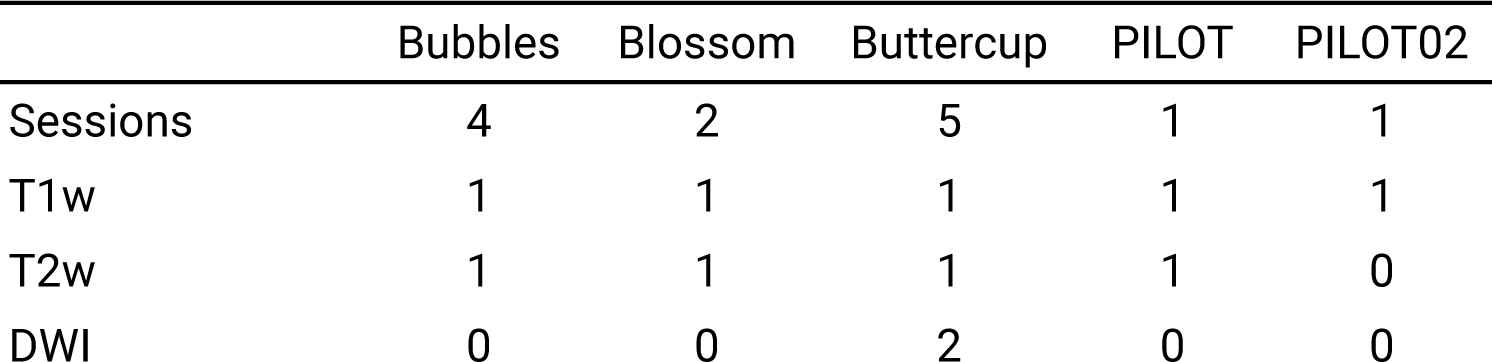
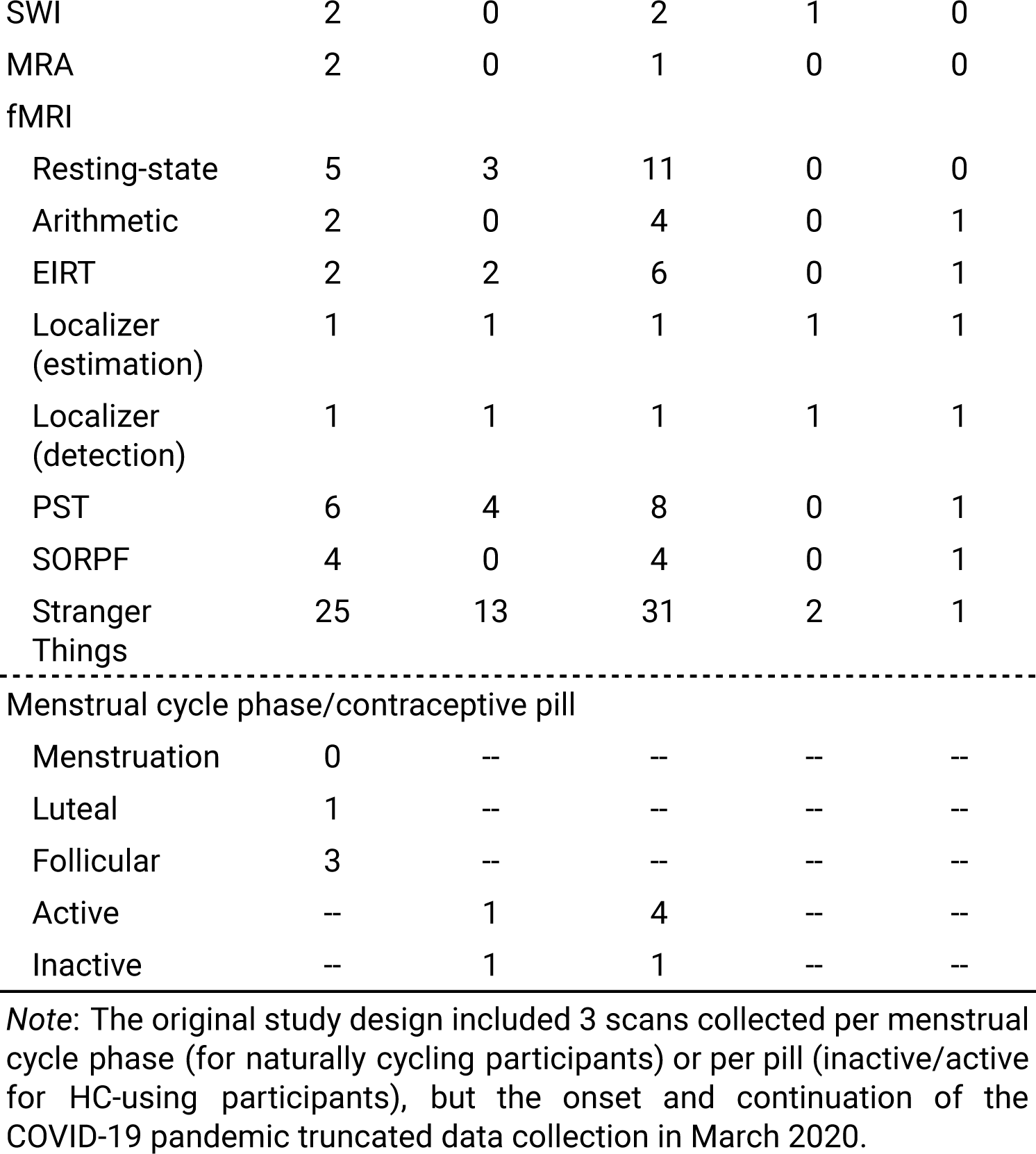
Number of runs per MRI sequence per participant.

|  | Bubbles | Blossom | Buttercup | PILOT | PILOT02 |
| --- | --- | --- | --- | --- | --- |
| Sessions | 4 | 2 | 5 | 1 | 1 |
| T1w | 1 | 1 | 1 | 1 | 1 |
| T2w | 1 | 1 | 1 | 1 | 0 |
| DWI | 0 | 0 | 2 | 0 | 0 |
| SWI | 2 | 0 | 2 | 1 | 0 |
| MRA | 2 | 0 | 1 | 0 | 0 |
| fMRI |  |  |  |  |  |
| Resting-state | 5 | 3 | 11 | 0 | 0 |
| Arithmetic | 2 | 0 | 4 | 0 | 1 |
| EIRT | 2 | 2 | 6 | 0 | 1 |
| Localizer<br>(estimation) | 1 | 1 | 1 | 1 | 1 |
| Localizer<br>(detection) | 1 | 1 | 1 | 1 | 1 |
| PST | 6 | 4 | 8 | 0 | 1 |
| SORPF | 4 | 0 | 4 | 0 | 1 |
| Stranger<br>Things | 25 | 13 | 31 | 2 | 1 |
| ----- |  |  |  |  |  |
| Menstrual cycle phase/contraceptive pill |  |  |  |  |  |
| Menstruation | 0 | -- | -- | -- | -- |
| Luteal | 1 | -- | -- | -- | -- |
| Follicular | 3 | -- | -- | -- | -- |
| Active | -- | 1 | 4 | -- | -- |
| Inactive | -- | 1 | 1 | -- | -- |

*Note:* The original study design included 3 scans collected per menstrual cycle phase (for naturally cycling participants) or per pill (inactive/active for HC-using participants), but the onset and continuation of the COVID-19 pandemic truncated data collection in March 2020.

### Participant information

**Location** participants.json, participants.tsv

**File format** javascript object, tab-separated values

Participants’ ages, responses to trait questionnaires, and hormonal contraceptive use are included; data have one line per participant.

**Location** sub-<subject>/<subject>.json, sub-<subject>/<subject>.tsv

**File format** javascript object, tab-separated values

All time-varying information, including responses to state questionnaires and activity tracking summaries, are included in tab-separated files named by participant. Data have one line per session/time point.

### MRI data

The number of each scan type and task acquired per participant are provided in Table 2.

#### Anatomical

**Location** sub-<subject>/ses-<session>/anat/sub-<subject>_ses-<session>_run-<run>_T1w.nii.gz, sub-<subject>/ses-<session>/anat/sub-<subject>_ses-<session>_run-<run>_T2w.nii.gz, sub-<subject>/ses-<session>/anat/sub-<subject>_ses-<session>_run-<run>_angio.nii.gz

**File format** NIfTI, gzip-compressed.

**Sequence protocols**

sub-<subject>/ses-<session>/anat/sub-<subject>_ses-<session>_run-<run>_T1w.json, sub-<subject>/ses-<session>/anat/sub-<subject>_ses-<session>_run-<run>_T2w.json, sub-<subject>/ses-<session>/anat/sub-<subject>_ses-<session>_run-<run>_angio.json

The defaced, raw, high-resolution anatomical images.

#### Fieldmap

**Location**

sub-<subject>/ses-<session>/fmap/sub-<subject>_ses-<session>_acq-<dwi/func>_dir-<AP/PA> _run-<run>_epi.nii.gz

**File format** NIfTI, gzip-compressed.

**Sequence protocols**

sub-<subject>/ses-<session>/fmap/sub-<subject>_ses-<session>_acq-<dwi/func>_dir-<AP/PA> _run-<run>_epi.json

The anterior-to-posterior and posterior-to-anterior fieldmaps for each diffusion-weighted and functional image collected.

#### Diffusion-weighted

**Location** sub-<subject>/ses-<session>/dwi/sub-<subject>_ses-<session>_run-<run>_dwi.nii.gz, sub-<subject>/ses-<session>/dwi/sub-<subject>_ses-<session>_run-<run>_dwi.bval, sub-<subject>/ses-<session>/dwi/sub-<subject>_ses-<session>_run-<run>_dwi.bvec

**File format** NIfTI, gzip-compressed.

**Sequence protocols**

sub-<subject>/ses-<session>/dwi/sub-<subject>_ses-<session>_run-<run>_dwi.json Diffusion-weighted images, along with files of b-values and -vectors.

#### Functional

**Location**

sub-<subject>/ses-<session>/func/sub-<subject>_ses-<session>_task-<task>_run-<run>_echo-<1-4>_part-<mag/phase>_bold.nii.gz,

sub-<subject>/ses-<session>/func/sub-<subject>_ses-<session>_task-<task>_run-<run>_echo-<1-4>_part-<mag/phase>_sbref.nii.gz,

**File format** NIfTI, gzip-compressed.

**Sequence protocols**

sub-<subject>/ses-<session>/func/sub-<subject>_ses-<session>_task-<task>_run-<run>_echo-<1-4>_part-<mag/phase>_bold.json,

sub-<subject>/ses-<session>/func/sub-<subject>_ses-<session>_task-<task>_run-<run>_echo-<1-4>_part-<mag/phase>_sbref.json, task-<task>_bold.json

**Participant responses**

sub-<subject>/ses-<session>/func/sub-<subject>_ses-<session>_task-<task>_run-<run>_events. tsv

Per-echo phase and magnitude images for each run of each task, along with single-band reference images.

The optimally combined data (i.e., one file per run) are available in `derivatives/fMRIPrep/.`

#### Susceptibility-weighted

**Location**

sub-<subject>/ses-<session>/swi/sub-<subject>_ses-<session>_acq-qsm_echo-<1-2>_part-<mag/phase>_coil-H<1-32>_GRE.nii.gz

**File format** NIfTI, gzip-compressed.

**Sequence protocols**

sub-<subject>/ses-<session>/swi/sub-<subject>_ses-<session>_acq-qsm_echo-<1-2>_part-<ma g/phase>_coil-H<1-32>_GRE.json

Per-coil, per-echo phase and magnitude data from susceptibility weighted scans.

#### Processed MRI data

All processed MRI data are provided in derivatives/ and organized according to the BIDS Standard.

#### Annotations

**Location** ses-<session>_task-strangerthings_acq-<annotator>_run-<run>_events.tsv

**File format** Tab separated values.

**Description** task-strangerthings_events.json

Annotations of each Stranger Things episode are included for each run.

#### Physiological recordings

**Location**

sub-<subject>/ses-<session>/func/sub-<subject>_ses-<session>_task-<task>_run-<run>_physio. tsv.gz

**File format** tab-separated values, gzip-compressed

**Acquisition, columns**

sub-<subject>/ses-<session>/func/sub-<subject>_ses-<session>_task-<task>_run-<run>_physio. json

Physiological recordings collected concurrently with functional scans. Filtered physiological recordings are provided in `derivatives/PhysioComb/…`.

## Technical Validation

To report the quality and characteristics of various data collected here, we provide the following data-specific metrics. For activity tracking, participants’ daily resting heart rate, number of active minutes, and hours of sleep are reported (Table 3). Averages, counts, and standard deviations (where applicable) of all self-report measures are presented in Tables 4 and 5. Salivary hormone concentrations per participant per measurement are summarized in Figure 2. Quality of scan-concurrent physiological recordings include heart rate (BPM), kurtosis, and heuristic quality (Zhao & Zhang, 2018) ECG data (Figure 3), average proportion of power in noise frequency bands (i.e., >0.5 Hz) for EDA recordings, and respiratory rate (breaths/minute) for respiration data. Quality of MR images varies by modality and includes IQMs calculated by MRIQC and QSIprep (Tables 6 - 9). For each task, we report average responses, response time, and accuracy, where appropriate (Tables 10 - 14). Naturalistic and resting state paradigms lack any cued responses. Finally, we summarize participants’ reported wakefulness throughout the scan and perceived effort on the fMRI tasks.

**Table 3.** Summaries of actigraphy data from wearable FitBit devices throughout data collection.

| Measure<br>(minutes) | Bubbles |  | Blossom |  | Buttercup |  |
| --- | --- | --- | --- | --- | --- | --- |
| | $\bar{x}$ | $\sigma$ | $\bar{x}$ | $\sigma$ | $\bar{x}$ | $\sigma$ |
| Sleep |  |  |  |  |  |  |
| Awake % | 13.5 | 2.1 | 12.8 | 2.5 | 15.8 | 3.1 |
| REM | 107.1 | 21.1 | 85.4 | 31.2 | 81.8 | 28.1 |
| Light | 233 | 40.4 | 267.7 | 56.2 | 232.8 | 55.1 |
| Deep | 81.8 | 16.7 | 84.2 | 13.6 | 68.0 | 23.7 |
| Exercise |  |  |  |  |  |  |
| Sedentary | 788 | 221.4 | 667 | 82.6 | 711 | 102.3 |
| Lightly Active | 220 | 87.0 | 225 | 52.3 | 154 | 47.5 |
| Fairly Active | 10 | 13.4 | 15 | 10.7 | 5 | 9.4 |
| Very Active | 20 | 41.1 | 14 | 16.8 | 2 | 4.5 |
| Missing days | 4 |  | 2 |  | 0 |  |
| Total days | 24 |  | 8 |  | 34 |  |

Note: Sleep data does not include daytime naps. All measurements are in minutes unless otherwise stated. Column names: $\bar{x}$ denotes sample mean; $\sigma$ , sample standard deviation.

**Table 4.** Average values of self-report trait measures across participants.

| Measure | $\bar{x}$ | $\sigma$ |
| --- | --- | --- |
| BIS/BAS |  |  |
| BAS Drive | 3.17 | 0.72 |
| BAS Fun Seeking | 3.25 | 0.62 |
| BAS Reward Responsiveness | 3.67 | 0.49 |
| BIS | 3.05 | 1.12 |
| Gender identity |  |  |
| Felt woman | 3.50 | 1.22 |
| Felt man | 0.20 | 0.45 |
| Felt both | 0.17 | 0.41 |
| Felt neither | 0.00 | 0.00 |
| Contentment with affirmed gender | 3.36 | 1.29 |
| Performing gender (W) | 0.67 | 1.00 |
| Performed gender (W) | 2.00 | 1.60 |
| Math Anxiety | 53.00 | 37.30 |

*Note:* Descriptive statistics presented are averaged across participants. *Felt woman/man/both/neither* scoring ranges from 0 to 5, *contentment* ranges from 0 to 5, and *performing/performed gender* is out of 2.

**Table 5.** Average values of self-report state measures across data collection.

| Scale | Bubbles | Blossom | Buttercup |
| --- | --- | --- | --- |

| | $\bar{x}$ | $\sigma$ | $\bar{x}$ | $\sigma$ | $\bar{x}$ | $\sigma$ |
| --- | --- | --- | --- | --- | --- | --- |
| PSQI | 4.50 | 1.69 | 5.50 | 1.00 | 12.2 | 1.55 |
| Duration | 0.38 | 0.74 | 0.25 | 0.50 | 0.50 | 0.53 |
| Disturbance | 0.88 | 0.35 | 1.00 | 0.00 | 2.00 | 0.00 |
| Latency | 0.63 | 0.52 | 2.75 | 0.50 | 2.90 | 0.32 |
| Day dysfunction | 0.75 | 0.46 | 0 | 0 | 1.8 | 0.63 |
| Sleep efficiency | 1.63 | 0.52 | 0.50 | 0.58 | 0.3 | 0.48 |
| Overall quality | 0.25 | 0.46 | 1 | 0 | 1.7 | 0.48 |
| Needs meds | 0 | 0 | 0 | 0 | 3 | 0 |
| PSS | 9.63 | 4.53 | 8.75 | 3.27 | 25.00 | 4.15 |
| Leisure Score Index | 47.63 | 10.72 | 42.75 | 3.90 | 8.10 | 2.70 |
| PANAS |  |  |  |  |  |  |
| Negative affect | 12.63 | 1.73 | 14.25 | 2.49 | 26.40 | 4.08 |
| Positive Affect | 41.75 | 4.58 | 44.25 | 4.82 | 17.90 | 3.14 |
| Fear | 7.50 | 1.00 | 9.50 | 1.50 | 17.80 | 2.64 |
| Hostility | 7.13 | 0.93 | 10.25 | 1.30 | 13.80 | 1.54 |
| Guilt | 6.00 | 0.00 | 6.00 | 0.00 | 15.80 | 5.13 |
| Sadness | 5.63 | 0.70 | 5.75 | 1.30 | 14.20 | 2.09 |
| Joviality | 33.00 | 4.39 | 33.75 | 3.27 | 13.20 | 2.09 |
| Self-assurance | 24.75 | 2.59 | 24.25 | 2.05 | 7.90 | 1.37 |
| Attentiveness | 16.25 | 2.05 | 16.75 | 1.64 | 7.20 | 2.09 |
| Shyness | 4.50 | 0.71 | 5.25 | 1.30 | 6.10 | 1.30 |
| Fatigue | 7.50 | 1.12 | 4.75 | 0.83 | 16.00 | 2.28 |
| Serenity | 9.50 | 0.71 | 10.50 | 0.87 | 3.90 | 1.04 |
| Surprise | 5.13 | 2.52 | 8.75 | 1.30 | 4.00 | 0.77 |

### Activity tracking

To inform the quality of activity tracking and its resulting data, we computed summaries of both sleep and exercise metrics. While not objective measures of quality, these summaries provide readers and potential users of this data with information concerning the consistency and similarity of data collected, within and between participants, respectively, across the study.

FitBIt data covered the period of active data collection for each participant, during which only Bubbles and Blossom had days with missing data. Daily summaries show average sleep duration was between 6.5 and 7.5 hours per night, similar to that of the average American (Jones, 2013). These summaries only include nightly sleep totals, not naps throughout the day (which are also recorded by FitBit actigraphy and included in the data). Exercise summaries show that participants spent the majority of their days sedentary, but with a range of lightly, fairly, and very active minutes.

### Self-report measures

Here we provide descriptive statistics for each of the self-report measures that participants completed over the course of the study. State measures were administered biweekly, on the same days as saliva sampling and MRI data collection. Trait measures were administered once, at the beginning of the initial MRI visit.

#### Trait measures

Because each trait measure was only administered once per participant, the responses are summarized across the study, to provide information to readers and potential users of these data about where these participants fall, in general, on each scale. Small standard deviations of BIS/BAS scores limit the study of inter-individual variability in behavior activation and inhibition. A large standard deviation of the math anxiety scores indicate a range of math-related anxiety is present in this sample, facilitating assessment of inter-individual differences in mathematical processing as a function of related anxiety. However, these data do not have sufficient power to make population-level claims about associations between these trait measures and any other neural or behavioral measure in this dataset. These data are best used in conjunction with other datasets (see Data Usage, Table 16). High *felt-woman*, *performing (woman) gender,* and *contentment with affirmed gender* scores on the Gender Identity scale, combined with low *felt-man, felt-both, and felt-neither* scores indicate that each of the participants are cis-gender women without gender dysphoria.

#### State measures

Self-report state measures are summarized in Table 5.

Total PSQI scores range from 0 (better) to 21 (worse), with scores greater than 5 indicative of poor overall sleep quality; less than 5, good sleep quality. Subscores of the PSQI range from 0 (better) to 3 (worse). These scores provide additional information about the participants’ sleeping habits and experience, to complement the FitBit sleep duration summaries. While the FitBit summaries are largely similar across participants, these measures indicate differences in experience across participants.

Perceived stress scores (PSS) range from 0 (lower) to 40 (higher) with scores less than 13 indicating low stress; 14 to 26, moderate stress; and greater than 27, high perceived stress. Two participants perceived low stress, while one perceived moderate stress, while all three participants’ standard deviations of perceived stress were similar. Relatively large standard deviations with respect to average values indicate intra-individual variability in addition to inter-individual variability.

The Godin-Shepard Leisure-Time Exercise Questionnaire’s Leisure Score Index (LSI) is a weighted combination of minutes spent engaged in light, moderate, and strenuous exercise. It has been validated against other scales and physiological measures and is widely used elsewhere in biomedical research (Amireault et al., 2015; Godin & Shephard, 1985). Higher scores indicate more time spent engaged in exercise and physical activity. Again, these scores align with the FitBit summaries of physical activity.

Per the post-scan debriefing questionnaires, one participant briefly fell asleep during the film watching task and another might have fallen asleep once during a scanning session. However, during those sessions both participants rate their effort on the tasks at 100% indicating that they were awake for at least some of the functional scans. Participant effort for each task in each session was 100%. The one exception was Buttercup’s effort on the arithmetic task in sessions 2 and 3, which she reported as 0%.

### Hormone data

Here, time points are delineated by “menstrual cycle day”, which in naturally cycling participants refers to the number of elapsed days since the onset of their most recent menses and in HC-using participants refers to the number of elapsed days since beginning their current 28-day pill pack (Figure 2). Overall, salivary estradiol and progesterone levels in the naturally cycling participant (Bubbles) roughly approximate expected trends throughout ovulation and the luteal phase (Figure 2, left, blue). Salivary estradiol levels in HC-using participants (Blossom, Buttercup) appear to do the same, though not enough data was collected from Blossom to characterize hormone trends. On the other hand, salivary progesterone levels in HC-using participants appear dampened throughout the menstrual cycle. This effect, unaltered estradiol and dampened progesterone, replicates that found in a similar study, 28andMe, which collected daily endocrine and MRI data from a single participant across one naturally-cycling menstrual cycle and one HC-using menstrual cycle (forthcoming). Cortisol levels vary between participants across the menstrual cycle, as well (Figure 2, right). While participants were instructed to collect saliva samples upon waking each day, the time of collection varied between participants across the study (Bubbles: 8:06 AM ± 6 minutes; Blossom: 6:51 AM ± 31 minutes; Buttercup: 10:47 AM ± 118 minutes).

### Physiological data

Despite the presence of MR-related artifacts in ECG recordings (detailed in (Bottenhorn et al., 2021)), the computed SQIs indicate that the data are acceptable, at least, even before denoising (Figure 3, top left) and that filtering modestly improved the quality of data.

Objective quality metrics are less established for measuring skin conductance via electrodermal activity (EDA). Fourier transformations were performed on each participant’s EDA recordings to provide information about the frequencies present (see example in Figure 1, right column).

Respiratory rate was calculated per participant per session, to assure that values fell within a normal range (i.e., 12 - 20 breaths per minute, (Chourpiliadis & Bhardwaj, 2022)). Average respiratory rates across scans across sessions were a little higher than the resting range (Table 6), which may be due to increased participant anxiety in the scanner.

**Table 6.** Average respiratory rate during fMRI scans per participant per session.

| Participant | Session | $\bar{x}$ | $\sigma$ |
| --- | --- | --- | --- |
| Blossom | 1 | 21.44 | 2.29 |
|  | 2 | 28.31 | 2.88 |
| Bubbles | 1 | 25.58 | 2.92 |
|  | 2 | 21.29 | 2.13 |
|  | 3 | 25.78 | 3.62 |
|  | 4 | 24.38 | 2.02 |
| Buttercup | 1 | 26.07 | 1.44 |
|  | 2 | 22.12 | 1.03 |
|  | 3 | 21.56 | 1.04 |
|  | 4 | 22.33 | 1.14 |
|  | 5 | 21.38 | 0.68 |

Note: PILOT and PILOT02 did not have physiological data collected during MRI sessions

### MRI data

#### MR image quality

Image quality metrics (IQMs) computed by MRIQC are presented in Tables 7 - 9 for T1-weighted, T2-weighted, and functional MRI data. For context, crowd-sourced data for each IQM is presented alongside our estimates (see *MRIQC Web API* columns in Tables 7 - 9), representing descriptive statistics aggregated across participants and sessions from a host of other datasets that have used MRIQC (Esteban, Blair, et al., 2017). Diffusion-weighted IQMs are presented in Table 10.

Structural image quality metrics (IQMs) are grouped into four categories: noise, information theory, specific artifacts, and other. Noise measurements include the *coefficient of joint variation, contrast-to-noise* ratio (CNR), *signal-to-noise* ratio (SNR), *Dietrich’s SNR*, and *Mortamet’s quality index 2* (QI2). The coefficient of joint variation between gray and white matter estimates the severity of head motion and intensity non-uniformity (INU) artifacts (Ganzetti et al., 2016), in which lower values indicate better image quality. The CNR between gray and white matter estimates tissue-type contrast (Magnotta et al., 2006), in which higher values indicate higher quality due to greater separation of gray and white matter. SNR is calculated for each tissue type (i.e., gray matter, white matter, cerebral spinal fluid) and the whole image, in which higher values indicate better image quality. Dietrich’s SNR calculates SNR using the air background for reference (Dietrich et al., 2007) and higher values indicate better image quality. Mortamet’s QI2 uses artifactual intensities in the air mask to calculate a goodness-of-fit Chi-square distribution (Mortamet et al., 2009), in which lower values indicate better image quality.

Information theory measurements include the *entropy-focus criterion* (ERC) and *foreground-background energy ratio* (FBER). The ERC estimates ghosting and blurring from head motion by calculating the Shannon entropy of intensities across voxels such that lower values indicate better image quality (Atkinson et al., 1997). The FBER compares the mean energy values from voxels inside the head to voxels outside the head (Shehzad et al., 2015), such that higher values indicate better image quality.

Assessments of specific artifacts included INU summary statistics, *Mortamet’s quality index 1* (QI1), and the *white matter to maximum intensity ratio*. INU summary statistics include the maximum, minimum, and median values of the bias field as calculated by N4ITK (Tustison et al., 2010), in which values closer to 1 indicate better image quality and values further from 0 indicate greater field inhomogeneity. Mortamet’s QI1 represents the proportion of voxels corrupted by artifacts, divided by the number of background voxels (Mortamet et al., 2009), in which lower values indicate better image quality. The white matter to maximum intensity ratio divides the median white matter intensity by the 95th percentile of the full image intensity, in which values between 0.6 and 0.8 indicate better image quality.

Other IQMs include the spatial smoothness of the image, the volume fractions of each tissue type, the residual partial volume effect for each tissue type, descriptive statistics for each tissue type, and the overlap of tissue probability maps (TPMs) with those of the ICBM nonlinear-asymmetric 2009c template. Spatial smoothness is calculated as the full-width at half-maximum (FWHM) of the voxel intensity distribution (Forman et al., 1995), in which lower values indicate better image quality and higher values indicate more blur. Volume fractions of each tissue type are based on total intracranial volume and should fall within a normal range (e.g., the distribution of volume fractions from the MRIQC Web API). Summary statistics including mean, standard deviation, and 90% confidence intervals were calculated for each tissue type and the image background. Finally, TPM overlap indicates the correspondence between each tissue type map and those of the ICBM template, in which higher values indicate better image quality.

Functional image IQMs from MRIQC include a number of spatial measures described above (i.e., EFC, FBER, smoothness, SNR, and summary statistics), in addition to temporal measures and artifact-specific measures (Table 9).Temporal measures include the temporal derivative of the root mean squared variance across voxels over time (DVARS), the *global correlation*, and temporal SNR. DVARS represents how the BOLD signal changes over the course of a functional acquisition (Power et al., 2012). Global correlation summarizes correlations between voxel time series across the brain (Saad et al., 2013). Temporal SNR represents the average BOLD signal over the course of a functional acquisition, divided by the standard deviation across the functional acquisition (Krüger & Glover, 2001), in which higher values indicate better image quality.

**Table 7.** T1-weighted image quality metrics compared with crowd-sourced values from MRQC’s web API.

| Metric | DIVA | MRIQC Web API |
| --- | --- | --- |
| Coefficient of joint variation | $0.31 \pm 0.02$ | $0.52 \pm 0.23$ |
| Contrast-to-noise ratio | $4.06 \pm 0.15$ | $2.84 \pm 0.87$ |
| Entropy-focus criterion | $0.46 \pm 0.02$ | $0.63 \pm 0.08$ |
| Foreground-background energy ratio | $24786.65 \pm 8935.60$ | $4935.91 \pm 20656.91$ |
| Smoothness (FWHM) | $4.36 \pm 0.24$ | $3.88 \pm 0.70$ |
| x | $4.4 \pm 0.21$ | $3.96 \pm 0.94$ |
| y | $4.55 \pm 0.26$ | $4.09 \pm 1.01$ |
| z | $4.13 \pm 0.27$ | $3.57 \pm 0.56$ |
| Volume fraction, CSF | $0.21 \pm 0.01$ | $0.2 \pm 0.04$ |
| Volume fraction, GM | $0.44 \pm 0.01$ | $0.43 \pm 0.04$ |
| Volume fraction, WM | $0.35 \pm 0.02$ | $0.37 \pm 0.02$ |
| Bias field, median | $0.5 \pm 0.06$ | $1.02 \pm 0.22$ |
| Bias field, range | $0.25 \pm 0.08$ | $0.42 \pm 0.17$ |
| Mortamet quality index 1 | $0.0 \pm 0.0$ | $0.01 \pm 0.02$ |
| Mortamet quality index 2 | $0.0 \pm 0.0$ | $0.03 \pm 0.1$ |
| Residual partial volume effect, CSF | $25.04 \pm 0.85$ | $26.42 \pm 7.78$ |
| Residual partial volume effect, GM | $12.6 \pm 0.6$ | $12.55 \pm 3.34$ |
| Residual partial volume effect, WM | $18.33 \pm 1.62$ | $16.75 \pm 4.43$ |
| Signal-to-noise ratio, CSF | $1.76 \pm 0.07$ | $2.4 \pm 0.89$ |
| Signal-to-noise ratio, GM | $9.61 \pm 0.63$ | $10.4 \pm 2.42$ |
| Signal-to-noise ratio, WM | $23.99 \pm 2.89$ | $18.25 \pm 4.01$ |
| Signal-to-noise ratio, Total | $11.79 \pm 0.94$ | $10.35 \pm 1.91$ |
| Dietrich's SNR, CSF | $25.37 \pm 2.63$ | $30.83 \pm 31.3$ |
| Dietrich's SNR, GM | $88.5 \pm 3.13$ | $59.87 \pm 56.28$ |
| Dietrich's SNR, WM | $132.84 \pm 4.78$ | $80.08 \pm 69.67$ |
| Dietrich's SNR, Total | $82.24 \pm 3.32$ | $56.92 \pm 52.13$ |
| WM to maximum intensity ratio | $0.89 \pm 0.02$ | $0.58 \pm 0.12$ |

**Table 8.**
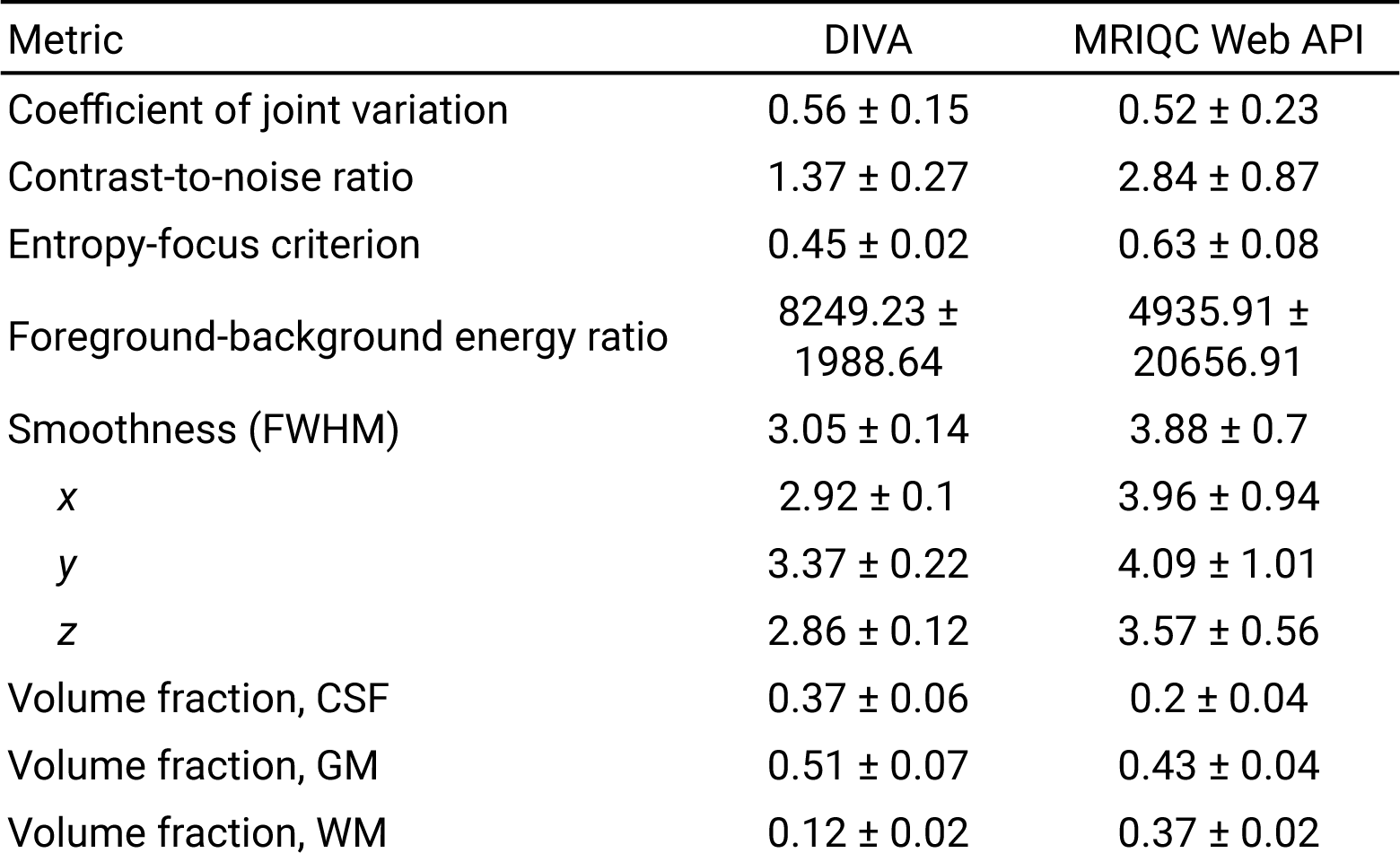
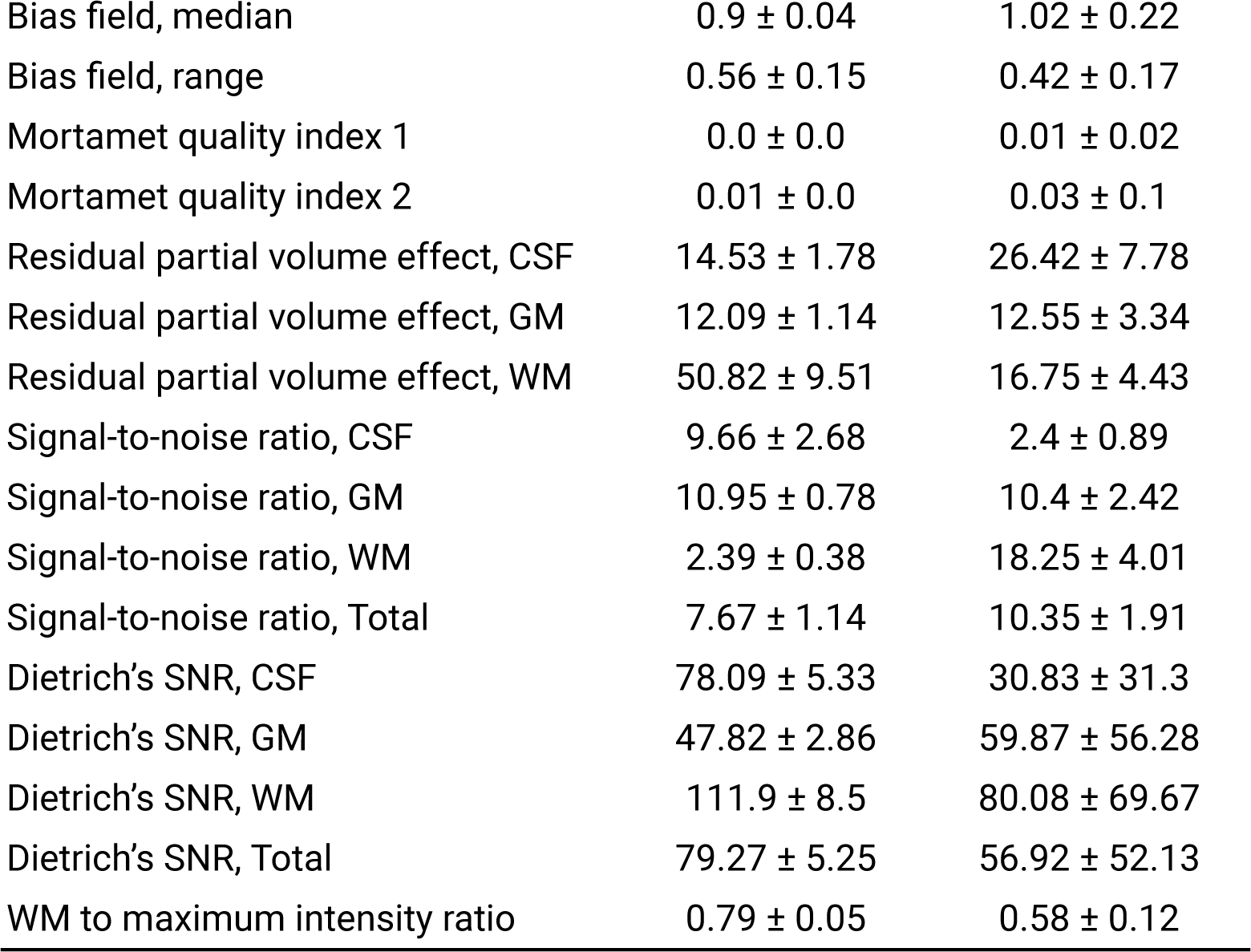
T2-weighted image quality metrics compared with crowd-sourced values from MRQC’s web API.

**Table 9.** BOLD functional image quality metrics compared with crowd-sourced values from MRQC’s web API.

| Metric | DIVA | MRQC Web API |
| --- | --- | --- |
| Spatial metrics |  |  |
| Signal-to-noise ratio | $2.71 \pm 0.30$ | $4.39 \pm 1.06$ |
| Entropy-focus criterion | $0.49 \pm 0.043$ | $0.49 \pm 0.063$ |
| Foreground-background energy ratio | $5976.77 \pm 2369.73$ | $4.64 \times 10^6 \pm 2.60 \times 10^7$ |
| Smoothness (FWHM) | $2.72 \pm 0.30$ | $2.65 \pm 0.30$ |
| $x$ | $2.42 \pm 0.19$ | $2.55 \pm 0.34$ |
| $y$ | $2.87 \pm 0.32$ | $2.99 \pm 0.39$ |
| $z$ | $2.85 \pm 0.46$ | $2.41 \pm 0.26$ |
| Temporal metrics |  |  |
| DVARS | $31.25 \pm 7.52$ | $28.14 \pm 9.59$ |
| Global correlation | $0.012 \pm 0.0075$ | $0.042 \pm 0.036$ |
| Temporal SNR | $48.94 \pm 14.69$ | $55.27 \pm 16.34$ |
| Artifacts, etc. |  |  |
| Framewise displacement | $0.11 \pm 0.025$ | $0.27 \pm 0.21$ |
| Ghost-to-signal ratio ($x, y$) | $-0.0078 \pm 0.013,$<br>$0.038 \pm 0.025$ | $-0.011 \pm 0.0094,$<br>$0.025 \pm 0.024$ |
| Outlier ratio (AFNI) | $0.0011 \pm 0.00090$ | $0.0049 \pm 0.0060$ |
| Quality index (AFNI) | $0.0046 \pm 0.0022$ | $0.0090 \pm 0.0059$ |
| Number of dummy scans | $0.30 \pm 0.61$ | $0.018 \pm 0.14$ |

Artifact-specific metrics include framewise displacement (FD), ghost-to-signal ratio (GSR), outlier ratio, quality index, and number of dummy scans. Framewise displacement quantifies head motion across the functional acquisition (Jenkinson et al., 2002; Power et al., 2012), including the average head motion across the acquisition, the number of frames above the FD threshold (0.2mm), and the percent of frames above the FD threshold; in all cases, lower values indicate higher quality. Ghost-to-signal ratio divides the intensity of the signal in the air space where ghosting is found along the phase-encoding axes by the intensity of the signal in the brain mask, such that lower values indicate better quality. Both outlier ratio and quality index are calculated by AFNI and represent the average proportion of outliers in each time point across each functional acquisition and the average Spearman’s correlation (i.e., 1 - *r_s_*) distance between each volume and the median volume, such that lower values for both measures indicate better quality. Finally, the number of dummy scans indicates the number of volumes identified as non-steady state at the beginning of each functional acquisition.

Increased neighboring DWI correlation (NDC) after processing with QSIPrep indicates a removal of noise and misaligned volumes, and a high NDC value (i.e., NDC > 0.7) aligns with high data quality ratings from expert reviewers in an independent study (Cieslak et al., 2021; Richie-Halford et al., 2022). Low maximum relative translation and no outlier slices indicate high-quality data, as well. Low mean framewise displacement (FD) indicates that the data are not likely corrupted by motion artifacts, although a maximum FD of 1.32 highlights the presence of some notable motion in the data. The T1w/DWI brain mask Dice distance indicates the dissimilarity of the b=0 mask from DWI data and a brain mask from the T1w scan such that 1 is perfect dissimilarity and 0 is perfect similarity. A low average value indicates that the T1w and DWI-computed brain masks exhibit low dissimilarity and, thus, a good deal of alignment and overlap.

### Task performance

#### Arithmetic task

Participant accuracy on the control trials was higher than on the math trials, and response time was lower compared with the mathematical conditions (Table 11). Overall, accuracy was relatively high, indicating that participants were, indeed, performing mathematical reasoning throughout the task. The average difficulty of mathematics trials, across conditions, was around 84% indicating that their difficulty did not exceed participants’ ability to solve the problems.

**Table 10.** DWI quality metrics from QSIPrep.

| Metric | Raw | Processed |
| --- | --- | --- |
| Neighboring DWI correlation | 0.766522 | 0.8116515 |
| Number of bad slices | 0 | 0 |
| Number of directions | 103 | 103 |
| Mean framewise displacement | 0.40320086 |  |
| Maximum framewise displacement | 1.3270673 |  |
| Maximum rotation | 0.00967422 |  |
| Maximum translation | 0.78596923 |  |
| Maximum relative rotation | 0.00609469 |  |
| Maximum relative translation | 0.57811721 |  |
| T1w/DWI brain mask Dice distance | 0.02244984 |  |

**Table 11.** Participant performance on the arithmetic task.

| Condition | Count | Accuracy | Response Time |
| --- | --- | --- | --- |
| Control | 56 | 95% | 0.85 ± 0.39 |
| Numeric, numeric | 7 | 100% | 0.84 ± 0.20 |
| Numeric, word | 12 | 92% | 0.79 ± 0.47 |
| Word, numeric | 15 | 100% | 0.68 ± 0.31 |
| Word, word | 22 | 91% | 1.09 ± 0.57 |
| Math | 112 | 84% | 1.02 ± 0.56 |
| Numeric, numeric | 37 | 81% | $0.89 \pm 0.47$ |
| Numeric, word | 28 | 86% | $1.04 \pm 0.41$ |
| Word, numeric | 25 | 84% | $1.04 \pm 0.55$ |
| Word, word | 22 | 91% | $1.12 \pm 0.82$ |

*Note:* Only Bubbles, Buttercup, and PILOT02 completed the arithmetic task. Within “Control” and “Math” conditions, labels refer to the representation of the equation and comparison, respectively.

An ANOVA found no significant difference in participant accuracy across trials with respect to condition (i.e., control vs. math), representation of the math problem (i.e., numeric or word), and representation of the solution (i.e., numeric or word). However, math trials showed slower response time than control trials (*F*(1,24) = 6.08, *p* = 0.02), as did trials using words to describe the answer, as opposed to numbers (*F*(1,24) = 4.70, *p* = 0.04).

#### Self-/Other-Referential Flanker Task

Mean response times and their standard deviations were similar across conditions (Table 12). Overall, participants found that more of the presented words did describe both themselves (Self) and the characters from Stranger Things (Other) than did not, and that nearly all of those descriptors were malleable characteristics (Control).

**Table 12.** Performance on the social conditions of the SORPF task across participants.

| Condition | # Trials | Response | Response Time |
| --- | --- | --- | --- |
| Self | 145 | $1.39 \pm 0.49$ | $0.53 \pm 0.25$ |
| Other | 146 | $1.44 \pm 0.50$ | $0.59 \pm 0.31$ |
| Control | 146 | $1.14 \pm 0.35$ | $0.61 \pm 0.32$ |

*Note:* Only Bubbles, Buttercup, and PILOT02 completed the SORPF task. There were only two choices for each trial, 1 (yes) or 2 (no).

Following a 2-way ANOVA, there were no differences in response time between congruent and incongruent Flanker trials (*F*(1,23) = 0.45, *p* = 0.51) or Flanker trials following self, other, and control conditions (*F*(2,23) = 0.30, *p* = 0.74), and there was no significant congruence by preceding condition interaction (*F*(2,23) = 0.07, *p* = 0.93). However, accuracy did differ between incongruent and congruent trials (*F*(1,23) = 1291.78, *p* < 0.001), though not between trials following self, other, and control conditions (*F*(2,23) = 0.57, *p* = 0.57), and with no significant congruence by preceding condition interaction (*F*(2,23) = 0.58, *p* = 0.57). These results are consistent with prior literature, which found no difference in response time during Flanker trials preceded by self-referential or control conditions (Alarcón et al., 2018).

#### Emotion Induction/Regulation Task

Few trials were missed indicating that participants were paying attention throughout the task. A lower average response for negative images than neutral indicates participants were paying attention to images and following directions. In line with prior research on this task (Blair et al., 2012), respenses (i.e., subjective ratings of an image’s valence) were significantly different between negative and neutral images for the viewing condition (*F*() = 349.61, *p* < 0.001) as were responses to negative images between the viewing and down-regulating conditions (*F*() = 83.98, *p* < 0.001). Reaction times, however, were not between negative and neutral images in the viewing condition (*F*() = 3.45, *p* = 0.06), but were between viewing and downregulating negative images (*F*() = 10.02, *p* = 0.0017). Together, these results indicate that the participants did, indeed, perceive the negative images more negatively than the neutral images and that downregulating that negativity was successful, with a slower response time during downregulation suggesting that slightly more effort was expended on these trials.

#### Probabilistic Selection Task

During the training runs, average participant accuracies were higher during the AB pair trials than during CD and EF pair trials, which corresponds with the proportion of “right” and “wrong” feedback given for correct responses for each of those trials (AB: 80/20, CD: 70/30, EF: 60/40). We observed ceiling effects for approach and avoidance reinforcement learning performance in the testing run. In the testing run participants chose the A stimulus on all Approach A trials (i.e., AC, AD, AE, AF stimuli pairs) and avoided the B stimulus on all Avoid B trials (i.e., BC, BD, BE, BF stimuli pairs). However, fMRI data from this task could still be used to assess Approach- and Avoidance-related processing during the testing phase. Additionally, perfect accuracy indicates successful learning during the training phase, data from which can be used to model trial to trial adaptation from average win-stay, loose-shift behavior during the training run (see *Data Usage* for more detail).

## Usage Notes

While the sampling scheme per participant was designed to maximize coverage of unique points across the menstrual cycle over the course of three months, day 24 was oversampled in one HC-using participant (Buttercup) and cycle/pill pack phases were not equally sampled across participants. This is due to a truncated experimental design due to the onset of the COVID-19 pandemic. However, this rich dataset has utility for several overarching reasons. Data collection was intended to contain similar measures to other neuroimaging datasets (Table 16), facilitating opportunities for multi-dataset integration. For example: ongoing work with DIVA data includes a transfer learning approach to studying contraceptive- and hormone-related resting-state functional connectivity with 28andMe and 28andOC data (Pritschet et al., 2020).

**Table 13.** Participant performance on the Flanker conditions of the SORPF task.

|  | # Trials | Accuracy | Response time |  |
| --- | --- | --- | --- | --- |
|  |  |  | Correct | Incorrect |
| Congruent | 131 | 100% | 0.52 ± 0.09 | n/a |
| Incongruent | 104 | 95% | 0.56 ± 0.69 | 0.43 ± 0.074 |

Note: Only Bubbles, Buttercup, and PILOT02 completed the SORPF task. There were no incorrect responses in the "Congruent" condition.

**Table 14.**
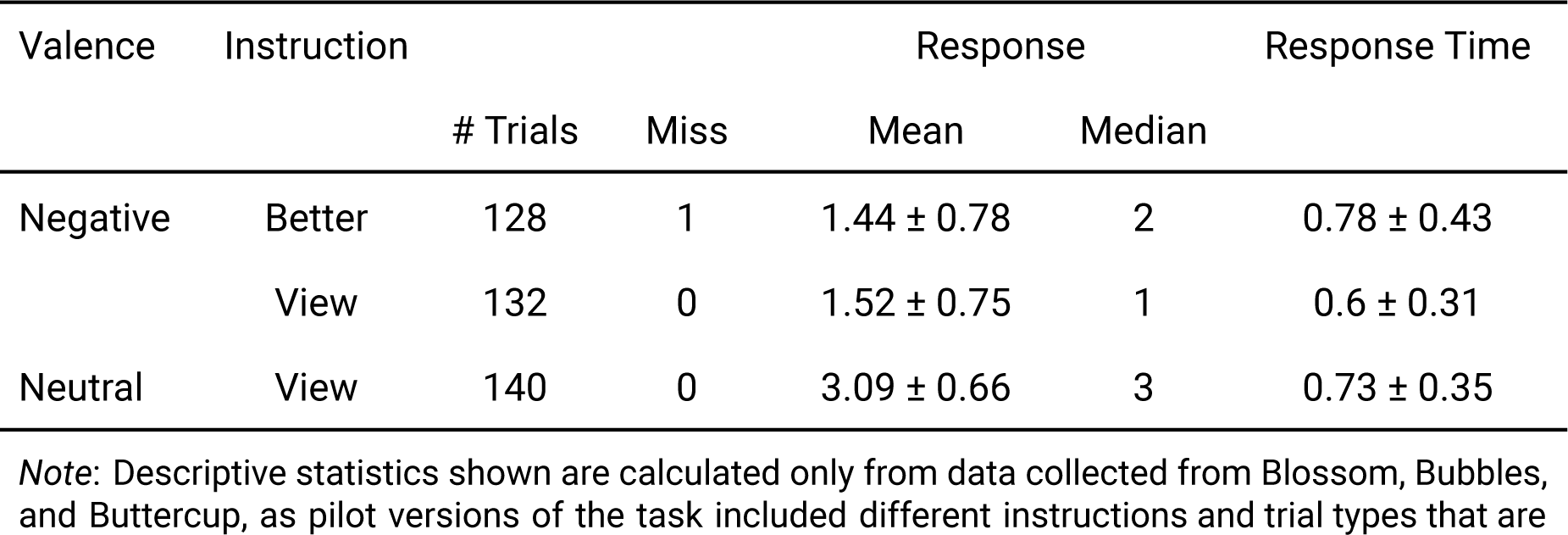
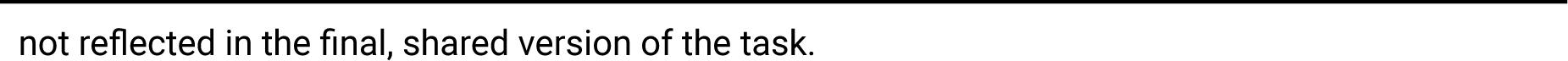
Descriptive characteristics of participant performance on the Emotion Induction/Regulation Task (EIRT), summarized across participants*.

| Valence | Instruction | Response |  |  |  | Response Time |
| --- | --- | --- | --- | --- | --- | --- |
|  |  | # Trials | Miss | Mean | Median |  |
| Negative | Better | 128 | 1 | 1.44 ± 0.78 | 2 | 0.78 ± 0.43 |
|  | View | 132 | 0 | 1.52 ± 0.75 | 1 | 0.6 ± 0.31 |
| Neutral | View | 140 | 0 | 3.09 ± 0.66 | 3 | 0.73 ± 0.35 |

Note: Descriptive statistics shown are calculated only from data collected from Blossom, Bubbles, and Buttercup, as pilot versions of the task included different instructions and trial types that are
not reflected in the final, shared version of the task.

**Table 15.** Participant performance on the Probabilistic Selection Task (PST)

| Condition | # trials per run | Accuracy | Response time |
| --- | --- | --- | --- |
| AB | 20* | $0.73 \pm 0.04$ | $0.90 \pm 0.19$ |
| CD | 20 | $0.57 \pm 0.18$ | $0.84 \pm 0.15$ |
| EF | 20 | $0.51 \pm 0.06$ | $0.85 \pm 0.13$ |
| Carrot | 40 | $1.00 \pm 0.00$ | $0.89 \pm 0.10$ |
| Stick | 40 | $1.00 \pm 0.00$ | $0.91 \pm 0.12$ |

*Note:* AB, CD, and EF represent stimulus pairs during the training run. “Carrot” and “stick” represent “choose A” and “avoid B” trials during the testing run. PILOT02 only had 18 trials for the AB pair during the training run. Reaction time is in seconds.

**Table 16.** Similarities between the DIVA dataset and other open and/or dense neuroimaging datasets.

| Dataset | Citation | Participants, sessions | Non-imaging data in common | Imaging data in common |
| --- | --- | --- | --- | --- |
| Midnight Scan Club | (Gordon et al., 2017) | 10, 12 | BIS/BAS | T1w, T2w, MRA, resting-state fMRI |
| Health Brain Network - Serial Scanning Initiative | (O'Connor et al., 2017) | 13, 14 | First day of last menstrual cycle, activity tracking | T1w, T2w, DWI, naturalistic film-viewing fMRI*, Flanker task fMRI (not SORPF), resting-state fMRI |
| Day2day | (Filevich et al., 2017) | 6, 43-50 | Salivary estradiol, activity tracking, caffeine and cigarette intake in the prior 24 hours, first day of last menstrual cycle, PANAS, sleep quality* | T1w, DTI, resting-state fMRI |
| MyConnectome | (Poldrack et al., 2015) | 1, 107 | Sleep quality*, PANAS, stress*, exercise* | T1w, T2w, DWI, resting-state fMRI |
| Individual Brain Charting | (Pinho et al., 2018) | 12, 8+ |  | T1w, T2w, DWI |
| Forrest Gump | (Hanke et al., 2014) | 37, up to 8 | Heart rate during scans*, respiration during scans | T1w, T2w, DTI, SWI, MRA, naturalistic film-viewing fMRI* |
| 28andMe, 28andOC | (Pritschet et al., 2020) | 1, 60 | PSS, mood*, estradiol* and progesterone*, HC use | T1w, resting-state fMRI |
| ABCD Study | (Casey et al., 2018) | 11800, 4+ | Exercise*, gender identity*, positive affect*, 24-hour caffeine and tobacco intake, HC use, salivary estradiol | T1w<sup>†</sup>, T2w<sup>†</sup>, DWI<sup>†</sup>, resting-state fMRI |
| Human Connectome Project | (Van Essen et al., 2012) | 1200, 1-2 | Affect*, stress*, menstrual cycle information, sleep quality | T1w, T2w, DWI, resting-state fMRI, naturalistic film-viewing fMRI<sup>§</sup>, somatosensory localizer fMRI* |

Note: \* A similar construct was assessed, but with a different instrument. <sup>†</sup> Same imaging sequence was used. <sup>§</sup> HCP film-viewing was collected at 7T.

### Self-report measures

The Gender Identity scores indicate, as previously mentioned, that all participants in this study, who were assigned female at birth, identify as cis-gender women and lack gender dysphoria.

Duplicate measures of sleep and exercise (i.e., from FitBit actigraphy and PSQI, Goldin) can be used to compare “objective” and “subjective” assessments of the same concept.

Post-scan debriefs include participants’ assessments of perceived task effort, wakefulness, and opinions about each major character in Stranger Things, per scanning session. Perceived task effort can be used as a quality metric for task-based fMRI data, as can wakefulness. Opinions about each major Stranger Things character, in terms of valence and arousal, can be used in conjunction with the SORPF task to investigate how emotional attachment to “Others” influences other-referential processing.

### Physiological data

The physiological data acquisition setup used here used an initial trigger pulse from the scanner was sent to the stimulus presentation computer, which then sent a signal to the BIOPAC acquisition module to indicate that a task was ongoing until turning it off at the end of the task run. In some fMRI runs, the trigger from the stimulus acquisition computer to the BIOPAC module did not fire. This issue is addressed in the shared data, but might result in slight timing differences for some scans. Based on these experiences, we recommend that future research sends trigger pulses per TR from the MRI scanner directly to the analog-to-digital converter (ADC) or other data acquisition (DAQ) device being used to collect peripheral physiological data.

Furthermore, sequence-specific MR-related artifacts were imparted on the ECG and EDA recordings, as mentioned above. These data have been preprocessed with PhysioComb (Bottenhorn et al., 2021) and are available in the `derivatives/` folder. Some heart rates are lower than the expected range (e.g., Bubbles, session 2), which is likely due to MR-related artifacts, but filtering out the MR-related noise generally increases estimated BPMs in this data (Bottenhorn et al., 2021). Researchers using these data should inspect heart rate before and after applying any filters to these data and consider incorporating SQIs to assess the impacts of their filtering strategy on ECG quality.

### MRI data

T1-, T2-, and diffusion-weighted data were acquired using the same sequence used by the ABCD Study, and thus the same metrics can be obtained using the ABCD processing pipeline (Hagler et al., 2019).

#### Multi-echo functional

Functional MRI data included in the DIVA dataset were acquired with a multiband, multiecho BOLD EPI sequence, with transverse acquisition for mitigating orbitofrontal signal dropout. Preprocessing of these data, including combining echoes to improve temporal SNR, is best done with fMRIPrep (Esteban et al., 2019) which includes tedana (DuPre et al., 2021) for multi-echo data processing. Furthermore, raw phase and magnitude images are provided for each echo of each functional scan. Phase data is commonly excluded from fMRI analyses, but can be used for phase regression or distortion correction and contains additional physiological information (e.g., respiratory and cardiac noise) (Petridou et al., 2009).

##### Naturalistic

Stranger Things was chosen as the naturalistic viewing used here because it is rich in socioaffective stimuli, representing a range of human interactions, emotional valence, and arousal. The episodes that participants watched during data collection were annotated by TR for visual and emotional information.

These data facilitate the study of both implicit, naturalistic emotion regulation (i.e., participants are instructed to remain still during the scans, prohibiting external emotional displays) and explicit, experimentally controlled emotion regulation (i.e., during EIRT). This is facilitated by the inclusion of valence and arousal in both the Stranger Things episode annotations and the use of IAPS images in the EIRT, supplemented with participant-rated emotional valence (i.e., 1 = most negative to 4 = most positive).

Furthermore, these data facilitate the study of implicit, naturalistic and explicit, experimentally controlled social processing by combining information from the post-scan debrief, SORPF task, and Stranger Things. The debrief includes participants’ emotional arousal and valence concerning each major character in Stranger Things. The SORPF task yields participant behavioral responses from the “Other” condition (i.e., participants were asked to view images of major Stranger Things characters and respond whether a given adjective described them) and measures of BOLD signal during this other-related social processing. Annotations from Stranger Things denote which major characters were on screen throughout a scan and the BOLD signal throughout these scans theoretically includes traces of character-specific social processing.

##### Tasks

The functional localizer task includes dissociable auditory, visual, and motor conditions for mapping of the corresponding primary sensory and motor regions.

The arithmetic task includes mathematical operations with both Arabic digits and number words, allowing for assessments of different elements of numerical cognition, which may be neurally dissociable (Skagenholt et al., 2018).

In the probabilistic selection task, participants’ choices can be used to evaluate whether they learned more from positive or negative feedback. Positive reinforcement learning performance is operationalized as the ability to choose stimulus A during testing, which has the highest probability of positive outcomes during training. Whereas negative reinforcement learning performance is operationalized as the ability to avoid choosing stimulus B during testing, which has the highest possibility of negative outcomes during training. Trial to trial adaptation can also be assessed as average win-stay, loose-shift behavior during the training run. Further participants’ choices in the training run can be analyzed using a Q-learning model (Frank et al., 2007; Frydecka et al., 2016). Specifically, individuals’ behavioral data can be quantitatively fit using separate learning-rate parameters for positive and negative feedback as done in prior work (Chase et al., 2010). An Empirical Bayes approach in which individual-level parameters are assumed to be sampled from a normally distributed population allows for computational algorithms to optimally estimate individual differences in learning parameters that are not directly observable in the data (Dombrovski et al., 2019).

#### Susceptibility-weighted

The susceptibility-weighted scans included in this dataset facilitate quantitative susceptibility mapping (QSM). Magnitude and phase images are included per echo per head coil channel (see *Data Records* for naming conventions). Phase reconstruction from multi-channel data can facilitate phase-offset and coil sensitivity corrections for improved image quality and accuracy (Haacke et al., 2015). QSM can be used to estimate iron content across the brain and to map venous blood and its oxygen saturation, providing complementary information to angiography in mapping brain vasculature.

#### Angiography

Information from MRA, SWI, and functional localizer fMRI scans can be used to study how the BOLD hemodynamic response varies across the brain with respect to arterial blood supply.

## Code Availability

All of the in-house scripts used to organize and process this data are available at https://github.com/NBCLab/diva-project, which also includes materials for each of the tasks detailed in this manuscript. External tools and packages are referenced and linked throughout.

## Acknowledgements

This project was supported by the Office of Research and Economic Development at Florida International University. We would like to thank Andrea Roman for her help with MRI sequence design and acquisition, Raquel Rodriguez for her help with data acquisition, Ulises Ley for his help annotating Stranger Things clips, Erica Musser for her guidance and assistance in setting up and acquiring physiological data, and Jenna Silva-Rosas for help with scheduling MRI data collection. Additional thanks to the FIU Instructional & Research Computing Center (**Error! Hyperlink reference not valid.** for providing the computing resources that contributed to the research results reported within this paper.

Special thanks to Dr. Essa Yacoub and the University of Minnesota Center for Magnetic Resonance Research (CMRR) for graciously providing access to the multiband EPI sequence used for data acquisition.

## Author Contributions (CRediT)

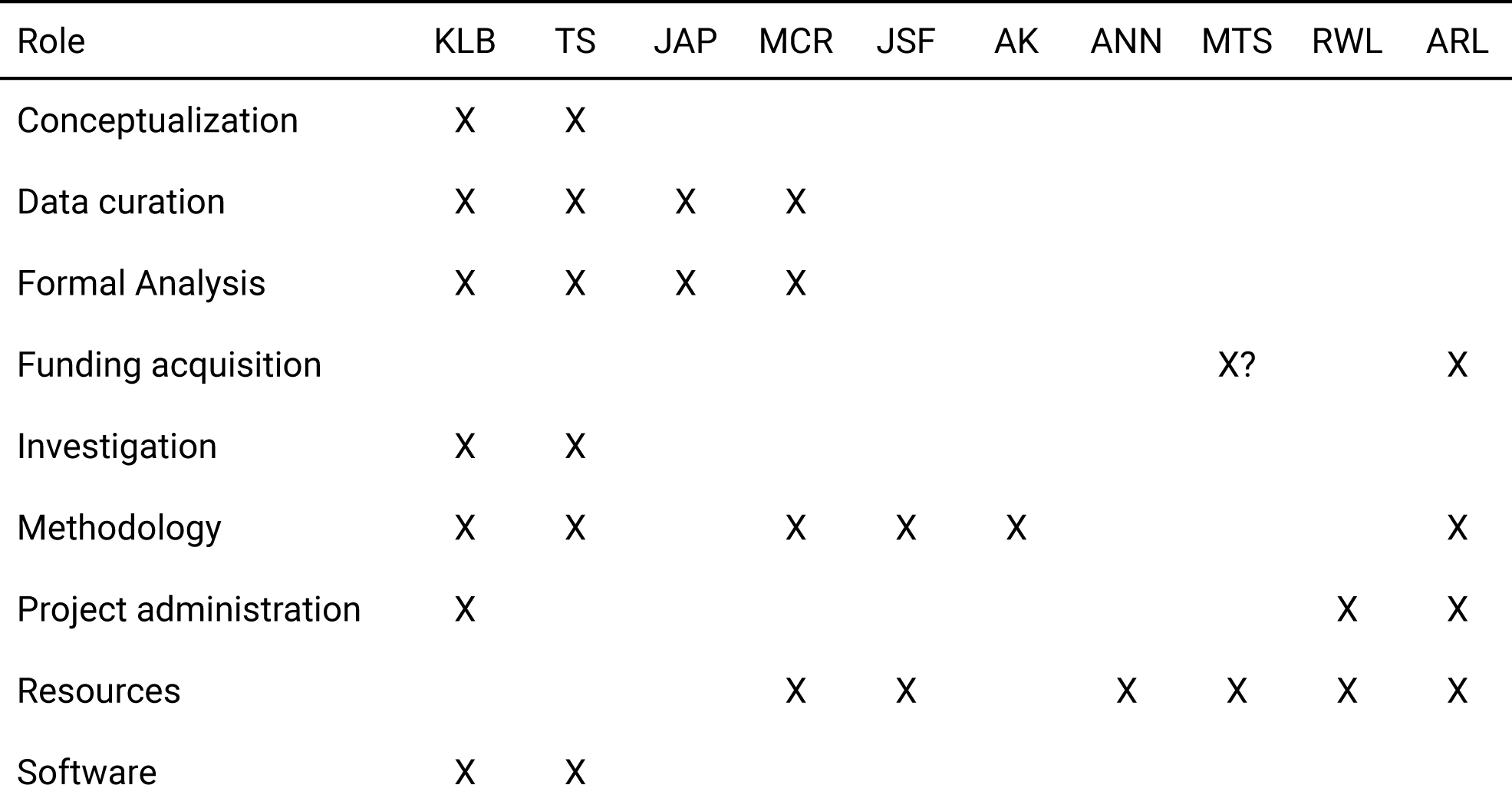
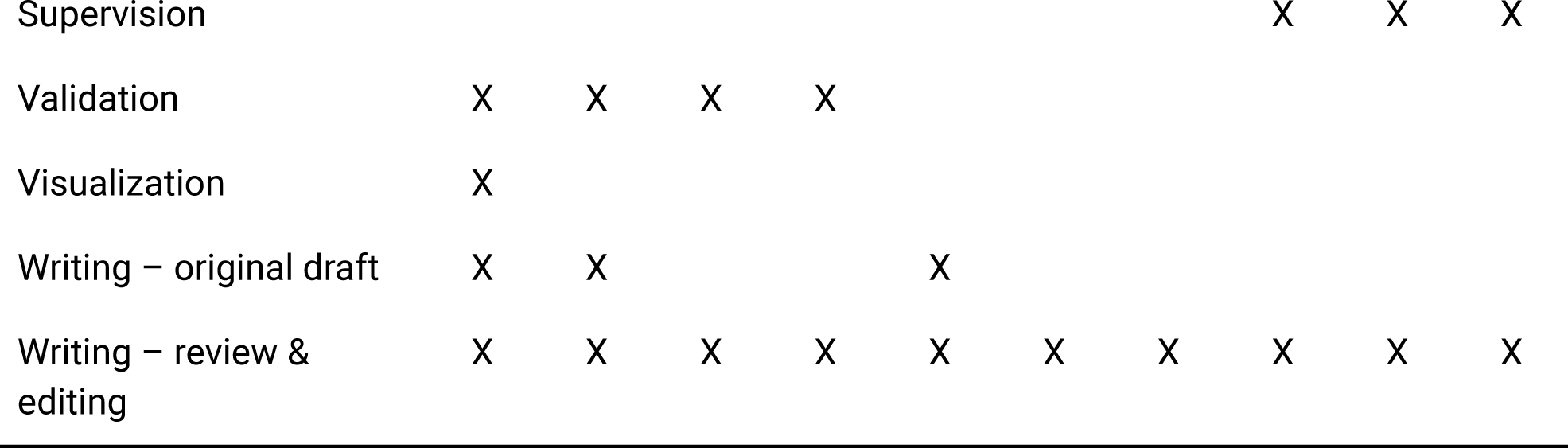

## Competing Interests

No authors have competing interests to declare.

## Notes

https://openneuro.org/datasets/ds002278

### Competing Interest Statement

The authors have declared no competing interest.

